# Catalytically inactive PARP1 protein drives PARP inhibitor–induced hematological toxicity

**DOI:** 10.64898/2026.09.21.753312

**Authors:** Xiaohui Lin, Zhengping Shao, Denitsa Yaneva, Wenxia Jiang, Demis Menolfi, Seema Khattri Bhandari, Brian J. Lee, Mirjam Schmucker, Faye Yan, Alan E. Tomkinson, Julian Stingele, Shan Zha

**Affiliations:** Institute for Cancer Genetics, Vagelos College of Physicians & Surgeons, Columbia University, New York, NY 10032, USA; Herbert Irving Comprehensive Cancer Center, Vagelos College of Physicians & Surgeons, Columbia University, New York, NY 10032, USA; Gene Center and Department of Biochemistry, Ludwig-Maximilians-Universität München, Feodor-Lynen-Str. 25, 81377 Munich, Germany; Departments of Internal Medicine and Molecular Genetics & Microbiology, and the University of New Mexico Comprehensive Cancer Center, University of New Mexico Health Sciences Center, Albuquerque, NM, 87131, USA; Cluster for Nucleic Acid Sciences and Technologies – NUCLEATE; Department of Pathology & Cell Biology, Vagelos College of Physicians & Surgeons, Columbia University, New York, NY 10032, USA; Department of Pediatrics, Vagelos College of Physicians & Surgeons, Columbia University, New York, NY 10032, USA; Department of Immunology & Microbiology, Vagelos College of Physicians & Surgeons, Columbia University, New York, NY 10032, USA

**Author notes:** Corresponding author: Dr. Shan Zha, Institute for Cancer Genetics, Vagelos College of Physicians & Surgeons, Columbia University; 1130 St. Nicholas Avenue, New York, NY 10032. Those authors made equal contributions (Xiaohui Lin, Zhengping Shao).

**Keywords:** PARP1, PARP inhibition, erythropoiesis, anemia, bone marrow failure

## Abstract

Dual PARP1/2 inhibitors (PARPi) selectively eliminate BRCA1/2-deficient cancers and represent the first targeted therapy for homologous recombination (HR)-deficient cancers. However, their use in maintenance therapy is limited by severe anemia and an increased risk for therapy-related leukemia. These toxicities are unexpected because PARP1 loss, which eliminates most DNA-damage-induced PARylation, does not cause anemia in mice. In contrast, PARP2 loss or catalytic inactivation causes anemia, motivating the development of PARP1-selective inhibitors. Using wild-type (WT), *Parp1^-/-^* and *Parp2^-/-^* mice, we show that hematopoietic toxicity of FDA-approved PARPi is driven primarily by inactive PARP1 rather than PARP2 inhibition. Accordingly, PARP1-selective inhibitors also cause PARP1-dependent anemia. Somatic expression of catalytically inactive Parp1 (*Parp1^E988A^*) causes lethal bone marrow failure, not found with somatic deletion of both *Parp1*&2. Mechanistically, inactive PARP1 obstructs the repair of diverse DNA lesions, including gaps, nicks, and Top1-cc, in contrast to the nick-selectivity of Parp2. In cells, inactive PARP1 compromises PARP2 recruitment to DNA lesions and causes severe genomic instability and mitotic bridges absent in Parp1&2-null cells. Thus, PARPi-induced hematopoietic toxicity is driven primarily by PARP1 inactivation, informing the design and use of next-generation PARP inhibitors.

## Introduction

DNA strand breaks activate PARP1 and PARP2, which use NAD+ to catalyze ADP-ribosylation of themselves and others (*e.g*., histones), thereby generating poly-ADP-ribose (PAR) chains (1). Although not essential for DNA repair, the highly negatively charged PAR chains promote chromatin relaxation and recruit PAR-binding DNA repair proteins (*e.g.*, XRCC1-LIG3) (2, 3). Dual PARP1/2 catalytic inhibitors (PARPi) selectively eliminate BRCA1- or BRCA2-deficient cells and have transformed the treatment for homologous recombination (HR)-deficient ovarian and breast cancers (4, 5). Over the past decade, four dual PARP1/2 inhibitors have received FDA approval (6, 7). However, maintenance use of two FDA-approved PARPi was suspended in 2022 because of severe anemia and a significantly increased risk of therapy-related myeloid leukemia (8, 9). These hematological toxicities are unexpected because complete loss of PARP1, which accounts for most DNA damage-induced PARylation, causes no overt hematopoietic defects in mice (10, 11). In contrast, PARP2-deficient mice develop mild yet reproducible anemia after 90 days of age (12), suggesting that PARP2 inhibition may contribute disproportionately to PARPi-associated hematologic toxicity. These and other findings have motivated the development of PARP1-selective inhibitors, including AZD5305 (saruparib), to improve hematologic safety while maintaining antitumor efficacy (13, 14). Accordingly, three of the four next-generation PARP inhibitors currently in clinical development are PARP1-selective.

While PARP1 and PARP2 are both activated by 5′-phosphorylated DNA ends exposed upon DNA strand break and share a similar catalytic mechanism, they differ in their catalytic robustness and lesion selectivity. PARP1, through its three N-terminal zinc-finger domains, recognizes and is activated by a broad spectrum of DNA lesions, including gaps, nicks (no missing base), TOP1 cleavage complexes (Top1cc), and DNA double-strand breaks (DSBs)(15–17). In contrast, PARP2 instead relies primarily on its WGR domain for DNA recognition, restricting its activation largely to 5′-phosphorylated nicks, the physiological substrates of DNA ligases (1, 2, 17–20). Because of its broader DNA lesion recognition and markedly higher cellular abundance, PARP1 is responsible for the vast majority of DNA damage-induced PARylation. Accordingly, deletion of PARP1, but not PARP2, significantly increases PARPi resistance in both cancer and normal cells (21).

In addition to inhibiting the enzymatic activity of PARP1 and PARP2, FDA-approved PARPi also extend the appearance of PARP1/2 proteins at the sites of DNA damage, a phenomenon referred to as “PARP trapping”(21–24). Under normal conditions, PARP1 and PARP2 are rapidly recruited to and activated by DNA lesions and undergo dynamic exchange, allowing each DNA break to activate multiple PARP molecules and thereby amplify the PARylation signal (15, 21, 24, 25). Mechanistically, however, the basis of PARP trapping differs between PARP1 and PARP2. While bulky NAD+ analogs (*e.g.*, BAD or EB47) allosterically retain PARP1 on DNA-ends *in vitro*, FDA-approved PARPi with significantly smaller footprints selectively induce allosteric retention of PARP2, not PARP1 (15, 21, 24, 25). Instead, the persistent accumulation of PARP1 at DNA damage sites primarily reflects its continuous recruitment to unrepaired DNA breaks resulting from delayed recruitment of the XRCC1-LIG3 complex, rather than direct allosteric stabilization of PARP1 on DNA(15, 21, 26, 27). Nevertheless, since PARP1/2 bind DNA breaks with nanomolar affinity (17, 18), if occupied the lesion, they can physically block downstream DNA repair enzymes (e.g., DNA LIG1/3 with micromolar affinity) and interfere with DNA repair. Thus, PARP trapping has been proposed as a major contributor to the therapeutic effect of PARPi in proliferating cancer cells (22, 27, 28). Correspondingly, we reported that in contrast to the normal development of *Parp2^-/-^* mice, mice expressing catalytically inactive *Parp2* (*Parp2^EA/EA^*) die of anemia by E16.5 because inactive PARP2 physically obstructs DNA Ligases I and III, delaying Okazaki fragment maturation in rapidly proliferating erythroblasts and phenocopying Lig1 deficiency (12, 29), further motivating the development of PARP1-selective inhibitors. Meanwhile, mice carrying a single catalytically inactive *Parp1* allele (*Parp1^+/E988A^*) died by E9.5, before fetal liver definitive hematopoiesis (30), raising the possibility that catalytically inactive PARP1 may also be toxic to erythroblasts even PARP1 loss is not.

Using *WT*, *Parp1^-/-^* or *Parp2^-/-^*mice treated with FDA-approved PARP inhibitor niraparib and Parp1-selective inhibitor saruparib and somatic expression of catalytical *Parp1^EA^*alone, we show that the hematologic toxicity of clinically approved PARP inhibitors is driven predominantly by catalytically inactive PARP1 rather than PARP2 or loss of PARP enzymatic activity. Using genetic mouse models, pharmacologic inhibition, and purified PARP1 protein on model DNA lesions, we demonstrate that inactive PARP1 binds to various DNA lesions, impeding their repair, whereas loss of PARP1 is well tolerated. Accordingly, cells expressing inactive Parp1 accumulate high levels of replication gaps, sister-chromatin exchanges, and mitotic breaks. These findings identify PARP1 trapping as the principal driver of hematologic toxicity and suggest that minimizing PARP1 trapping, rather than increasing PARP1 selectivity, as a strategy to minimize hematological toxicities.

## Results

### Niraparib-induced hematopoietic defects depend on PARP1, not PARP2

To investigate the hematologic toxicity of PARP inhibition *in vivo*, we administered niraparib to young adult mice by oral gavage once daily for five consecutive days. Dose-escalation studies identified 60 mg/kg/day as the optimal dose to reproducibly induce hematologic toxicity in *WT* (*Parp1^+/+^Parp2^+/+^*) mice. This dose corresponds approximately to the FDA-approved clinical dose of 300 mg/day for a 60-kg adult when converted using standard body surface area normalization (31, 32). Because red blood cells (RBC) in mice have a half-life of approximately 40 days, analyses at 5 days after the initial treatment did not affect peripheral RBC counts (Supplemental Figure 1A) and minimized the secondary systemic effects. Meanwhile, niraparib treatment decreased the peripheral white blood cell (WBC) counts by 50% in 5 days, with reductions in both short-lived neutrophils and newly generated lymphocytes (Figure 1A).

**Figure 1.**
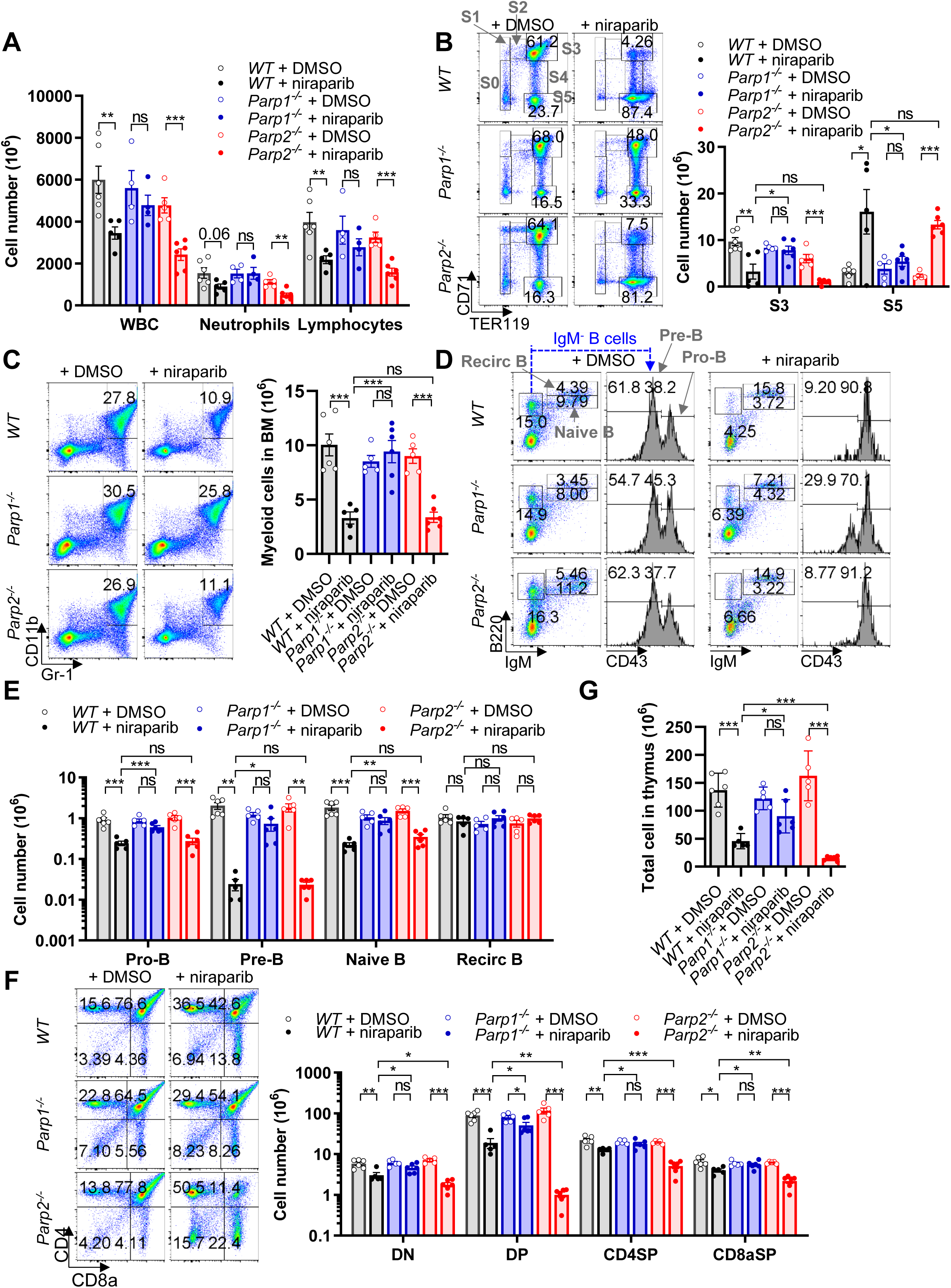
Deletion of PARP1, but not PARP2, rescues the niraparib-induced acute hematopoietic toxicity. (A) White blood cell (WBC), neutrophil and lymphocyte counts in peripheral blood from *WT*, *Parp1^-/-^* and *Parp2^-/-^* mice treated with niraparib. (B) Representative flow cytometry analysis and absolute numbers of the different developmental stages (S0-S5) of erythropoiesis in the BM from *WT*, *Parp1^-/-^* and *Parp2^-/-^* mice treated with niraparib. CD71 and TER119 staining were used to determine erythropoiesis stages in B220^-^Thy1.2^-^Gr-1^-^CD11b^-^ live cells. The defined subsets include S0 (CD71^Low^TER119^Low^), S1 (CD71^High^TER119^Low^), S2 (CD71^High^TER119^Mid^), S3 (CD71^High^TER119^High^), S4 (CD71^Mid^TER119^High^), S5 (CD71^Low^TER119^High^). (C) Representative flow cytometry analysis and absolute number of myeloid cells (CD11b^+^Gr-1^+^) in the BM from *WT*, *Parp1^-/-^* and *Parp2^-/-^* mice treated with niraparib. (D) Representative flow cytometry analysis and (E) absolute numbers of Pro-B cells (B220^+^IgM^-^CD43^+^), pre-B cells (B220^+^IgM^-^CD43^-^), naïve-B cells (B220^low^IgM^+^) and recirculate B cells (B220^high^IgM^+^) in the BM from *WT*, *Parp1^-/-^* and *Parp2^-/-^* mice treated with niraparib. (F) Representative flow cytometry analysis and absolute numbers of T cells at different developmental stages in the thymus from *WT*, *Parp1^-/-^* and *Parp2^- /-^* mice treated with niraparib. Defined subsets include double negative (DN; CD4⁻CD8a⁻), double positive (DP; CD4⁺CD8a⁺), CD4 single positive (CD4SP; CD4⁺CD8a⁻), and CD8a single positive (CD8aSP; CD4⁻CD8a⁺) T cells. (G) Absolute number of thymic cells in the thymus from *WT*, *Parp1^-/-^* and *Parp2^-/-^*mice treated with niraparib.

To understand how niraparib affects de novo hematopoiesis, we performed flow cytometry analysis of bone marrow (BM). The analyses revealed profound impairment of de novo erythropoiesis, reducing proliferating erythroblast counts (Ter119⁺CD71^high^, S3) by ∼80% while mature RBC (Ter119⁺CD71⁻, S5) accumulated (Figure 1B). Consistent with a 50% reduction of WBC, BM myeloid cells (CD11b^+^Gr-1^+^) also reduced by ∼50%, whereas BM B cells (B220^+^) declined by >80% (Figure 1C and Supplemental Figure1B). Consistent with the replication-associated toxicity of PARPi, within the B-cell compartment, the most pronounced effect was observed in highly proliferative large pre-B cells (B220⁺IgM⁻CD43⁻), which undergo clonal expansion following successful IgH rearrangement. This was accompanied by an almost complete depletion of the immediately downstream quiescent small pre-B cells, which undergo IgL rearrangement, and an ∼90% reduction in naïve B cells (B220^low^IgM⁺) (Figure 1D-E). In contrast, upstream pro-B cells (B220⁺IgM⁻CD43⁺) were relatively preserved, resulting in a dramatic decrease in the pre-B/pro-B ratio upon PARPi treatment (Supplemental Figure1C). Meanwhile, the quiescent recirculating mature B cells (B220^high^IgM^+^) were largely unaffected, leading to an increase in the recirculating/naïve B cell ratio (Figure 1E and Supplemental Figure1D). A similar pattern was observed in the thymus. CD4⁺CD8⁺ double-positive (DP) thymocytes, which actively undergo V(D)J recombination at the TCRα locus after clonal expansion triggered by productive TCRβ rearrangement, were preferentially depleted by niraparib, whereas mature single-positive thymocytes were relatively resistant (Figure 1F). Since DP thymocytes account for approximately 75% of total thymocytes, niraparib treatment resulted in a marked reduction in both thymic cellularity and thymus weight (Figure 1G and Supplemental Figure 1E–F). Together, these findings indicate that niraparib preferentially ablates proliferating hematopoietic progenitors, particularly erythroblasts and developing lymphocytes undergoing programmed clonal expansion. Consequently, their immediate downstream developmental populations, S3 erythroblasts, small pre-B and naïve B cells in the bone marrow, and DP thymocytes in the thymus, are preferentially depleted following niraparib treatment.

To address whether inactivation of PARP1 or PARP2 accounts for the hematological toxicity, we repeated these experiments in *Parp1^-/-^* and *Parp2^-/-^* mice (n>=4 for each genotype). Strikingly, deletion of Parp1 almost completely rescued niraparib-induced depletion of peripheral WBCs, bone marrow erythroblasts, myeloid cells, developing B cells, and thymocytes (Figure 1A–G and Supplemental Figure 1A–F). In contrast, Parp2 deficiency did not significantly rescue. Instead, niraparib caused even greater depletion of *Parp2^−/−^* thymocytes, particularly DP cells, than in WT controls (Figure 1F–G and Supplemental Figure 1E–F). Moreover, hematopoietic stem and progenitor cells (HSPCs), including LSK (Lin⁻Sca1⁺c-Kit⁺) and LK (Lin⁻Sca1⁻c-Kit⁺) populations, were also reduced by niraparib in *WT* and *Parp2^⁻/⁻^* mice while preserved in *Parp1^⁻/⁻^* mice, contributing to the decline of granulocytes in BM and the decline of neutrophils in peripheral blood (Supplemental Figure 1G–H). Together, these results demonstrate that niraparib-induced hematopoietic toxicity requires the presence of PARP1 but not PARP2, indicating that inactive PARP1, rather than loss of PARP catalytic activity or inactive PARP2, is the principal driver of PARP inhibitor–associated hematologic toxicity *in vivo*.

### Somatic expression of catalytically inactive PARP1 causes lethal bone marrow failure

To understand whether the hematological toxicity reflects a unique property of niraparib or a general consequence of the presence of inactive Parp1 protein, we turned to genetic models. Since *Parp1^+/E988A^* mice cannot transmit the E988A allele via germline (30), we generated a conditional E988A allele (*Parp1^AN^*) in which an FRT-NeoR-FRT cassette blocks Parp1-E988A transcription (30). While *Parp1^+/AN^* mice are normal and fertile, *Parp1^AN/AN^* mice succumbed in utero, likely due to leaky Parp1-E988A expression (30). We therefore created a conditional Parp1 allele (*Parp1^flox^*) by flanking exon 4 with LoxP sites (Supplemental Figure 2A–B). Crossing these alleles into *Rosa26a^FLP/CreER-T2^*mice (33), which express FLP constitutively together with a tamoxifen-inducible Cre recombinase (Cre-ERT2), enabled tamoxifen-mediated deletion of exon 4 and generated cells that express only the inactive Parp1-E988A in *Rosa26a^Flp/^*^CreER-T2^*;Parp1^AN/flox^* mice (hereafter *Parp1^EA/f^*) (Supplemental Figure 2A–C). To distinguish the effects of inactive PARP1 from combined loss of Parp1 and Parp2 together, we also generated *Rosa26a^+/^*^CreER-T2^*;Parp1/2^flox/flox^* mice (*Parp1/2^f/f^*) alongside Ctrl *Rosa26a^+/^*^CreER-T2^*;Parp1^+/flox^* and *Rosa26a^+/^*^CreER-T2^ mice using the previously characterized *Parp2^f/f^* allele generously provided by Dr. Françoise Dantzer (34). Oral gavage of tamoxifen (5 mg daily ×2) induced efficient recombination of *Parp1^flox^* and *Parp2^flox^* alleles and expression of *Parp1^E988A^*, as confirmed by PCR (Supplemental Figure 2C-D) and validated by Western blot for the absence of Parp1/2 proteins and DNA damage– induced PARylation (Supplemental Figure 2E).

Strikingly, all *Parp1^f/EA^* mice died within 13 days of tamoxifen treatment, whereas none of the *Parp1/2^f/f^* mice succumbed (Figure 2A). By day 10, *Parp1^f/EA^* mice exhibited a 20–25% (∼5 g) loss of body weight, compared to ∼5% (∼1 g) in *Parp1/2^f/f^* controls (Supplemental Figure2F). Peripheral blood analyses revealed an ∼80% reduction of all nucleated cell types in tamoxifen-treated *Parp1^f/EA^* mice, versus ∼40% in treated *Parp1/2^f/f^* mice (Figure 2B on a log scale). Total peripheral RBC counts were only moderately reduced by day 10, consistent with their ∼40-day half-life (Supplemental Figure 2G). Meanwhile, reticulocytes decreased by ∼60% in both treated genotypes (Figure 2C). Platelets with a 2–3 day half-life decreased by ∼50% in treated *Parp1^f/EA^*mice, but increased in treated *Parp1/2^f/f^* mice, potentially reflecting a compensatory feedback response (Figure 2D). Histological analyses revealed marked depletion of nucleated cells, including megakaryocytes, the precursor of platelets, in the BM of treated *Parp1^EA/f^*mice (Figure 2E-F), consistent with severe thrombocytopenia. Histological analyses also revealed diffuse pink staining in BM, consistent with hemolysis and invasion of fat cells in treated *Parp1^EA/f^* mice, but not *Parp1/2^f/f^*mice (Supplemental Figure 2H). In contrast, treated *Parp1/2^f/f^* mice showed a moderately reduced cellularity without hematolysis and a moderate increase of megakaryocytes (Figure 2D-F).

**Figure 2.**
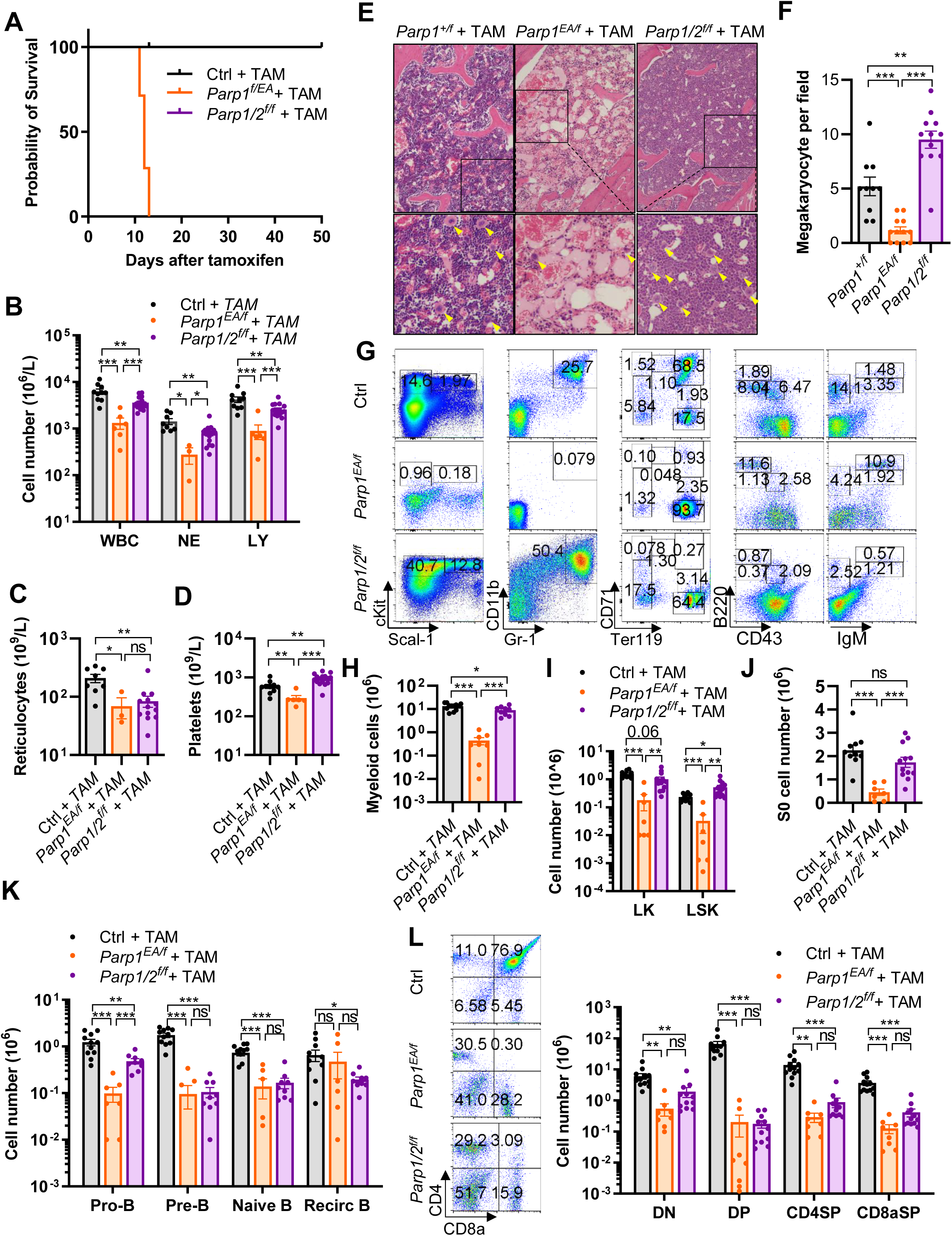
Expression of PARylation-deficient PARP1 solely somatically, but not loss of PARP1/2, leads to bone marrow failure. (A) Survival curves of *Rosa26a^Flp/Cre-ERT2^;Parp1^f/EA^*, *Rosa26a^+/Cre-ERT2^;Parp1/2^f/f^* and control mice following tamoxifen gavage. (B) WBC, neutrophil (NE) and lymphocyte (LY) counts, (C) reticulocyte counts, and (D) platelet counts in the peripheral blood from *Rosa26a^Flp/Cre-ERT2^;Parp1^f/EA^*, *Rosa26a^+/Cre-ERT2^;Parp1/2^f/f^* and control mice at day 10 following tamoxifen gavage. (E) Representative H&E stained BM sections from *Rosa26a^Flp/Cre-ERT2^;Parp1^+/f^*, *Rosa26a^Flp/Cre-ERT2^;Parp1^EA/f^*and *Rosa26a^+/Cre-ERT2^;Parp1^f/f^;Parp2^f/f^*mice following tamoxifen gavage. The upper panels show the 100× images, with boxed regions indicating fields used for (F) megakaryocyte quantification. The lower panels show higher-magnification images of the boxed regions, with yellow arrows indicating megakaryocytes. (G) Representative flow cytometry analysis and absolute numbers of (H) myeloid cells, (I) LK and LSK cells, (J) S0 erythroid cells, and (K) pro-B, pre-B, naïve B and recirculating B cells in the BM from mice of the indicated genotypes. (L) Representative flow cytometry analysis and absolute numbers of T cells at different developmental stages in thymus from mice of the indicated genotypes.

Moreover, BM, thymus, and spleen cellularity all decreased in treated *Parp1^EA/f^* mice. Specifically, by day 10 after TAM gavage, GR1^+^CD11b^+^ granulocytes were > 95% depleted in treated *Parp1^EA/f^* mice versus ∼30% depleted in treated *Parp1/2^f/f^*mice (Figure 2G-H). Correspondingly, HSPCs, including LSK and LK subsets, were decreased by ∼85% in treated *Parp1^EA/f^* mice (Figure 2I). In treated *Parp1/2^f/f^* mice, the LSK cells decreased by ∼30%, and the LK cells actually increased by ∼100%, consistent with potential feedback compensation (Figure 2I). Consistent with HSPCs depletion, early erythroid progenitors (CD71^Low^Ter119^Low^; S0), the earliest B cell progenitors (B220^+^IgM^-^CD43^+^; pro B cells), and T cell progenitors (CD4^-^CD8a^-^ double negative T cells) were also decreased by more than 4-fold in treated *Parp1^EA/f^* mice (Figure 2G and 2I-L). In contrast, treated *Parp1/2^f/f^* mice retained substantial populations of S0 erythroid progenitors, pro-B cells, and DN T cells in the BM and thymus (Figure 2J-L). Although the severe bone marrow failure and depletion of HSPCs limits our ability to analyses stage specific blockade within lymphoid lineage in treated *Parp1^EA/f^* and *Parp1/2^f/f^* mice, the ratio of naïve B vs Pre-B and SP vs DP T cells, which reflects V(D)J recombination at the IgL and TCRa loci, was significantly increased (Supplemental Figure 2I-J), reflecting quiescent mature cells are more resistant to inactive PARP1. V(D)J recombination itself is not affected by inactive PARP1(35). Meanwhile, in treated *Parp1^EA/f^*mice, peripheral splenic lymphocytes - B cells (B220^+^IgM^+^) and T cells (CD4^+^CD8a^-^ and CD4^-^CD8a^+^) - decreased by ∼80% and ∼65%, respectively, while remaining unchanged in treated *Parp1/2^f/f^* mice (Supplemental Figure 2K-N). Together, these results indicate that most severe PARPi-induced hematopoietic toxicity is an on-target consequence of inactivating PARP1 protein, not loss of both PARP1 and PARP2 or PARylation.

### The selective PARP1 inhibitor saruparib also causes PARP1-dependent anemia

Compared with dual PARP1/2 inhibitors, the PARP1-selective inhibitor saruparib exhibits substantially reduced GI toxicity; however, hematological toxicity is not completely eliminated, especially at high therapeutic doses (14). To determine whether this residual toxicity is likewise mediated by PARP1, we treated *WT* and *Parp1^-/-^* mice with 12 mg/kg/day saruparib, corresponding approximately to the FDA-approved clinical dose of 60 mg/day based on body surface area normalization(31). This dose reproducibly induced hematopoietic toxicity in WT mice after five consecutive daily doses and was therefore used for all subsequent analyses. Five days of saruparib reduced peripheral WBC counts by ∼40% in comparison to 50% by narparib (Figure 1A), affecting both neutrophils and lymphocytes, whereas peripheral RBC and platelet counts remained unchanged (Figure 3A and Supplemental Figure3A–B). Similar to niraparib, saruparib impaired erythropoiesis, reducing proliferating S3 erythroblasts by approximately 40% and resulting in a relative accumulation of mature S5 erythrocytes in BM (Figure 3B and Supplemental Figure 3C).

**Figure 3.**
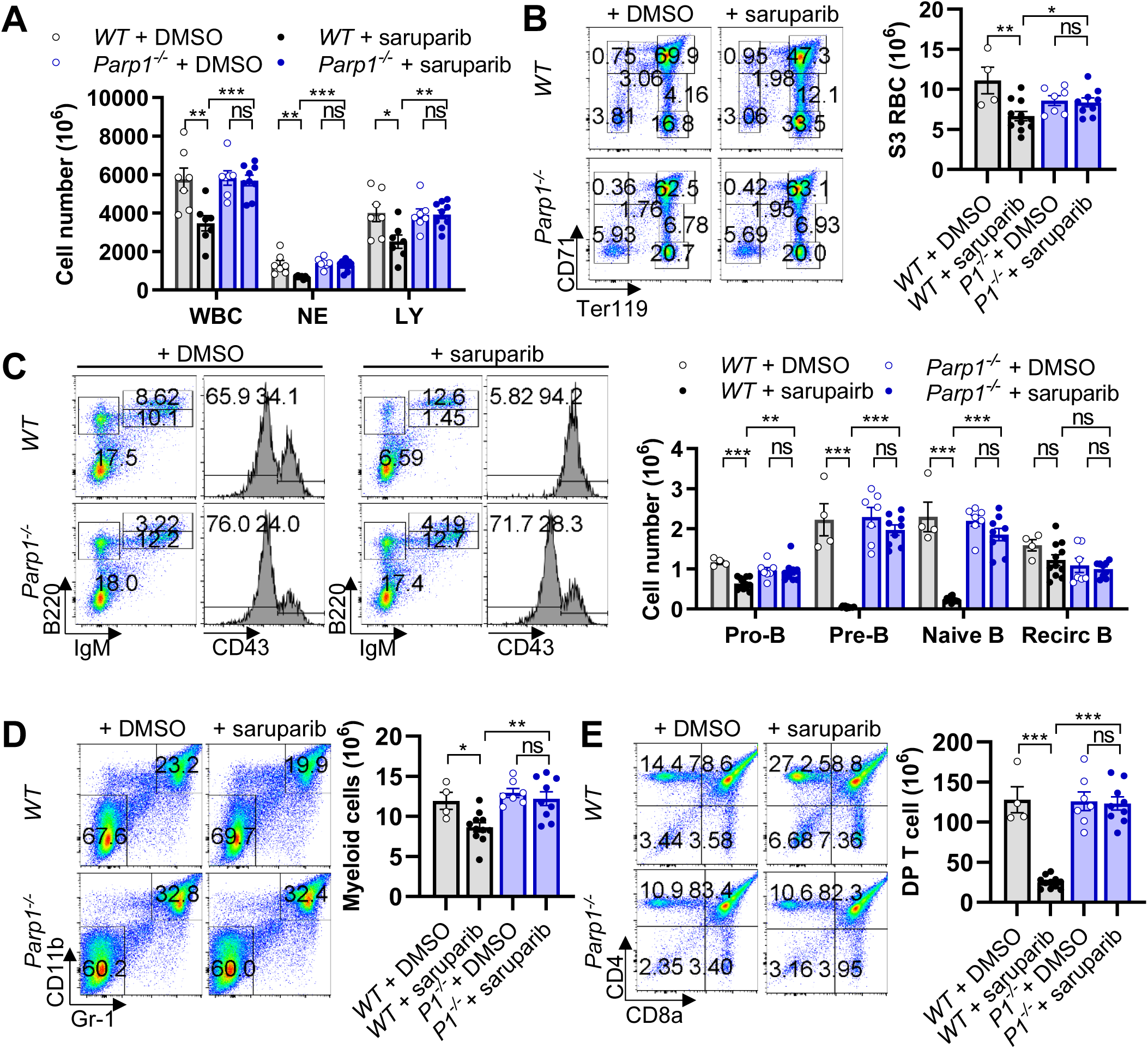
PARP1 inhibition underlies the hematopoietic toxicity induced by saruparib. (A) White blood cell, neutrophil and lymphocyte counts in peripheral blood from *WT* and *Parp1^-/-^* mice treated with saruparib. (B) Representative flow cytometry analysis of different developmental stages of erythropoiesis and absolute number of S3 erythoid cells in the BM from *WT* and *Parp1^-/-^*mice treated with saruparib. (C) Representative flow cytometry analysis and absolute numbers of pro-B, pre-B, naïve B and recirculate B cells in the BM from *WT* and *Parp1^-/-^* mice treated with saruparib. (D) Representative flow cytometry analysis and absolute number of myeloid cells in the BM from *WT* and *Parp1^-/-^* mice treated with saruparib. (E) Representative flow cytometry analysis of different developmental stages of T cells and absolute number of DP T cells in the thymus from *WT* and *Parp1^-/-^* mice treated with saruparib.

Within the B cell lineage, pre-B cells and naïve B cells after programmed clonal expansion were preferentially deleted, exhibiting greater than 90% decrease, accompanied by a marked reduction in the pre-B/pro-B ratio and corresponding increases in the naïve B/pre-B and recirculating B/naïve B ratios (Figure 3C and Supplemental Figure 3D–F). Bone marrow myeloid cells were also modestly reduced (∼10% vs 20% for niraparib) (Figure 3D). Likewise, in the thymus, DP thymocytes were preferentially depleted, resulting in a significant reduction in thymic cellularity and thymus weight (Figure 3E and Supplemental Figure 3G–K). Strikingly, deletion of Parp1 almost completely abolished saruparib-induced depletion of hematopoietic cell populations, including erythroid, myeloid, B-cell, and T-cell lineages (Figure 3 and Supplemental Figure 3). These findings demonstrate that a PARP1-selective inhibitor still causes hematological toxicity *in vivo,* and this toxicity also depends on the presence of inactive PARP1.

### Purified PARP1 protein blocks the repair of diverse DNA lesions in a catalytic activity-dependent manner

Unlike PARP2, which is selectively activated by 5′-phosphorylated nicks (without a missing base), PARP1 has high affinity for diverse DNA lesions. We therefore hypothesized that catalytically inactive PARP1 may interfere with multiple DNA repair reactions by persistently occupying DNA repair intermediates. To test this hypothesis, we reconstituted a series of *in vitro* enzymatic assays using purified recombinant PARP1-WT and catalytically inactive PARP1-E988A (PARP1-EA) proteins together with defined DNA substrates (Figure 4 and Supplemental Figure 4A–B). As observed for PARP2 (29), inactive (minus NAD+) PARP1 protein also inhibited LIG1-mediated 5′-phosphorylated DNA nick ligations in a dose-dependent manner (Figure 4A–B). Moreover, pharmacological inhibition of PARP1 (PARP1-WT + NAD^+^ + olaparib) or expression of catalytically inactive PARP1-EA (±NAD^+^) markedly inhibited T4 DNA ligase–mediated ligation of 5′-phosphorylated DNA nicks (Figure 4C-D, and Supplemental Figure 4), consistent with physical blockade rather than specific inhibition of DNA LIG1. Moreover, inactive PARP1 also blocked, terminal deoxynucleotidyl transferase (TdT)-mediated extension from 3′-OH DNA gaps (Figure 4E-F, and Supplemental Figure 5), and tyrosyl-DNA phosphodiesterase 1 (TDP1)-mediated removal of 3′-phosphotyrosyl adducts from nicked DNA substrates (Figure 4G-H, and Supplemental Figure 6). These inhibitory effects were observed with catalytically inactive PARP1 generated either genetically (PARP1-EA) or pharmacologically (PARP1-WT plus olaparib), demonstrating that persistent binding of inactive PARP1 sterically impedes the access of multiple DNA repair enzymes to their substrates. Together, these results support a model in which the presence of inactive PARP1 protein interferes with the processing of a broad range of DNA repair intermediates – from nick to polymerase filling and TDP1. This capacity to obstruct broader DNA repair provides a mechanism for why inactive PARP1, rather than inactive PARP2, is the predominant driver of PARP inhibitor–induced hematopoietic toxicity *in vivo*.

**Figure 4.**
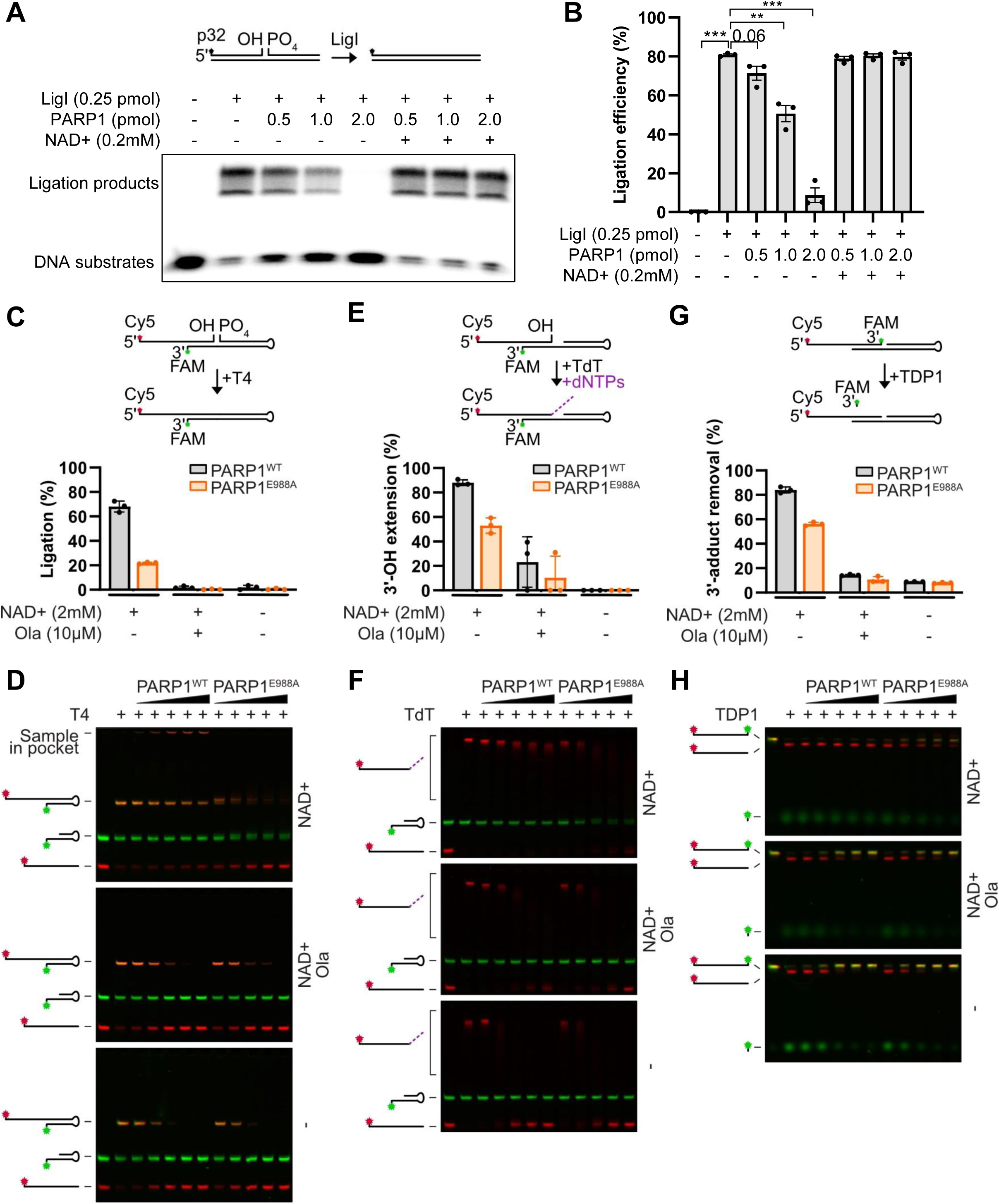
Inactive PARP1 blocks multiple DNA repair processes. (A) Representative gel image and (B) quantification of *in vitro* LIG1-mediated nick ligation in the presence of purified PARP1 protein, with or without NAD+. Schematics, quantifications, and representative images of UREA-PAGE gels showing the impact of PARP1 inhibition or inactivation on (C-D) T4-mediated 5′-phosphorylated nick ligation, (E-F) TdT–mediated 3′-OH gap DNA synthesis, and (G-H) TDP1–mediated hydrolysis of 3′-phosphodiester-linked FAM adducts in the presence of increasing amounts of recombinant PARP1 (WT or catalytically inactive E988A), with or without NAD⁺ and olaparib, as indicated. Values represent the means ± SD of three independent experiments for the highest PARP1 (WT or catalytically inactive E988A) concentration.

### The presence of Inactive PARP1, but not the absence of PARP1 causes severe genomic instability

To investigate the physiological consequences of inactive PARP1 in cells, we generated conditional embryonic stem (ES) cells and v-Abl–transformed pre-B (v-Abl B) cells from *Rosa26^Flp/Cre-ERT2^;Parp1^EA/f^*and control *Rosa26^Flp/Cre-ERT2^;Parp1^+/f^* mice (33,34). Treatment with 4-hydroxytamoxifen (4-OHT) converted the conditional Parp1^f^ allele to a null allele and generated *Parp1^EA/-^* cells expressing only one copy of catalytically inactive PARP1 and control *Parp1^+/-^* cells. Parallel experiments were performed in conditional *Rosa26^Cre-ERT2/+^;Parp1/2^f/f^*cells to compare the consequences of inactive PARP1 with complete loss of PARP1 and PARP2. Expression of PARP1-E988A as the sole PARP1 protein completely abolished ES cell proliferation (Supplemental Figure7A). Similarly, *Parp1^EA/-^* Abelson cells failed to proliferate following 4-OHT treatment (Figure 5A). Cell cycle analysis demonstrated a marked loss of BrdU-positive S-phase cells accompanied by prominent G1 arrest and accumulation of sub-G1 cells, consistent with cell-cycle arrest and apoptosis (Figure 5A and Supplemental Figure 7B). In contrast, deletion of both Parp1 and Parp2 produced a substantially milder proliferative defect (Figure 5A). To determine whether the toxicity of inactive PARP1 primarily resulted from replication-associated lesions, cells were treated with a CDK4/6 inhibitor to prevent entry into S phase before 4OHT treatments. Cell cycle inhibition significantly rescued viability of *Parp1^EA/-^*Abelson cells (Figure 5B, from 4.5% to 26%), consistent with replication-associated toxicity of PARP1 inhibition. We next examined DNA replication fork progression efficiency using DNA fiber analysis. While olaparib increased fork speed as previously documented (36), both *Parp1^EA/-^* and *Parp1/2^-/-^* cells show significantly slower fork progression (Figure 5C). Interestingly, replication forks progressed modestly faster in *Parp1^EA/-^* cells than in *Parp1/2^-/-^* cells, consistent with previous reports that PARP inhibition can paradoxically accelerate fork progression through PRIMPOL-dependent repriming (37–39). Moreover, the newly synthesized DNA fiber in *Parp1^EA/-^* cells was sensitive to S1 nuclease digestion, indicating the accumulation of single-stranded DNA gaps behind replication forks, (Figure 5C). These gaps are consistent with defective maturation of replication intermediates, including unligated Okazaki fragments and PRIMPOL-mediated repriming events previously observed following PARP inhibition (37–39).

**Figure 5.**
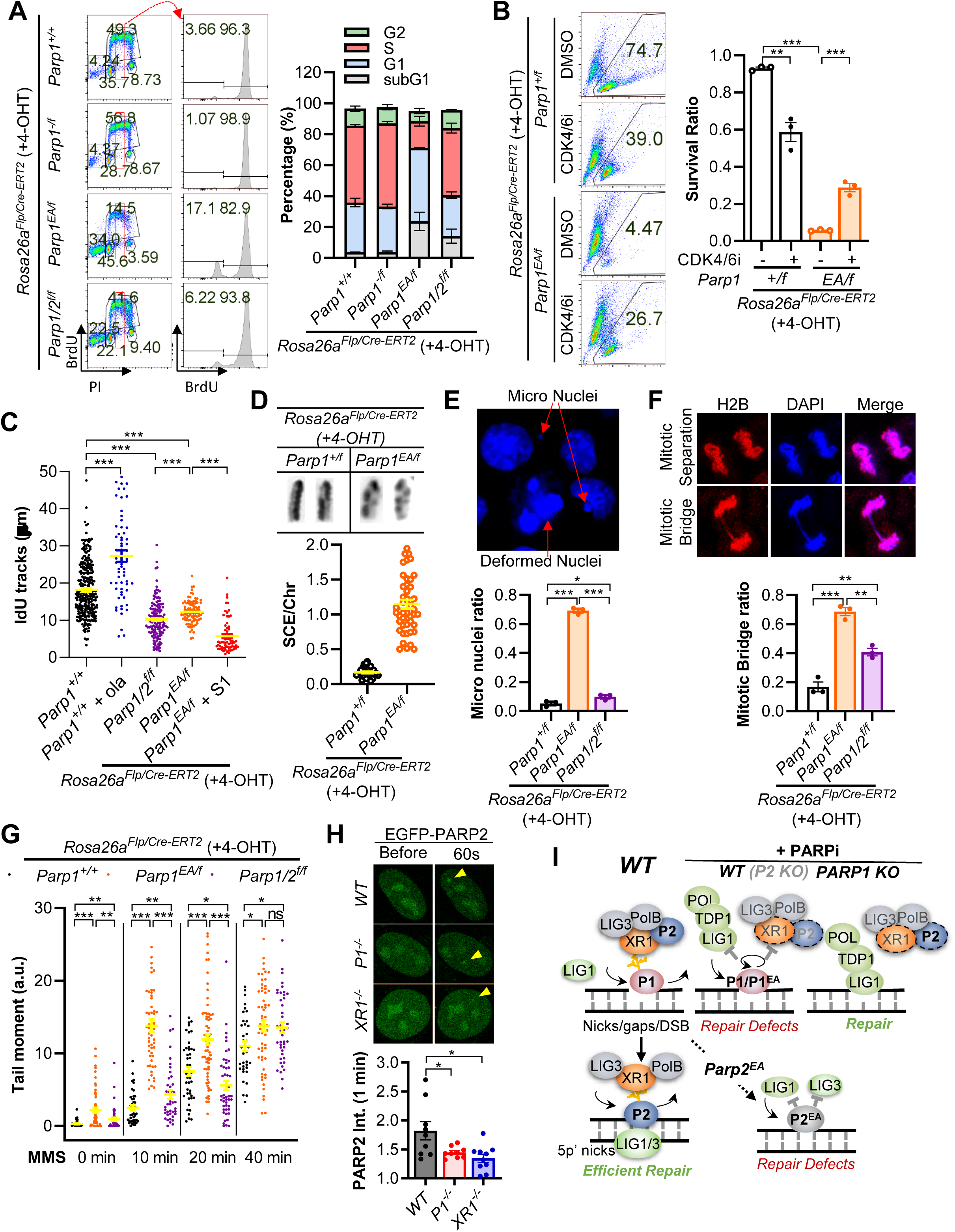
Inactive PARP1 causes more severe genomic instability than PARP1/2 loss. (A) Representative flow cytometry analysis and quantification of the cell cycle of v-Abl B cells of the indicated genotypes following 4-OHT treatment. (B) Representative flow cytometry analysis and quantification of survival ratios of v-Abl B cells of the indicated genotypes following 4-OHT treatment ± CDK4/6 inhibitor (PD-0332991, 1µM). (C) IdU track lengths show the replication fork speed in v-Abl B cells of indicated genotypes following 4-OHT treatment. Cells were pulse-labeled with 25 µM CldU for 10 min and then with 200 µM IdU for 20 min. At least 60 fibers per sample were quantified. Representative images and quantification of (D) sister chromatid exchange, (E) micro nuclei and (F) mitotic bridge in ES cells of the indicated genotypes following 4-OHT treatment. (G) Alkaline comet tail moment in total genomic DNA of v-Abl B cells of the indicated genotypes following 4-OHT treatment. Cells were treated with Methyl Methanesulfonate (MMS, 0.1 mg/mL) for indicated time before harvest. (H) Representative images and quantification of GFP-PARP2 recruitments upon micro irradiation in RPE-1 cells. (I) Working model: Under physiological conditions, PARP1 acts as the primary sensor of a broad spectrum of DNA lesions, including single-strand breaks, gaps, and double-strand breaks. Upon binding to damaged DNA, PARP1 catalyzes PAR synthesis, which recruits downstream repair factors, including XRCC1, LIG3, and Polβ. PARP1 also promotes the recruitment of PARP2, which preferentially recognizes 5′-phosphorylated DNA nicks generated during the final steps of DNA repair. Together, PARP1 and PARP2 coordinate efficient completion of DNA repair. Upon PARPi treatment, PARP1 becomes trapped on multiple types of DNA lesions. Trapped PARP1 sterically blocks the access of both the canonical base excision repair machinery (XRCC1/LIG3/Polβ) and backup repair pathways (polymerases, TDP1, and LIG1) to damaged DNA. In addition, PARP1 trapping prevents efficient recruitment of PARP2, further compromising DNA repair. Consequently, DNA repair defects in PARPi treated WT cells are primarily driven by trapped PARP1. Consistent with this model, genetic deletion of PARP2 has minimal additional impact on PARPi induced hematological toxicity because PARP2 recruitment is already impaired by PARP1 trapping. In contrast, deletion of PARP1 eliminates the physical barrier at DNA lesions, allowing downstream repair factors to access damaged DNA and partially restore repair, despite the absence of PARP dependent signaling. In genetically inactive PARP models, the mechanisms differ from pharmacological inhibition. Catalytically inactive PARP1 remains bound to a broad range of DNA lesions, thereby obstructing multiple DNA repair pathways. In contrast, catalytically inactive PARP2 is efficiently recruited by active PARP1 to 5 ′-phosphorylated DNA nicks, where it selectively blocks the final ligation step of DNA repair. Thus, although both inactive PARP1 and inactive PARP2 impair DNA repair, inactive PARP1 disrupts a broader range of repair processes and consequently causes more severe genomic instability and hematological defects than inactive PARP2.

Consistent with elevated replication stress, *Parp1^EA/-^*cells exhibited widespread genomic instability, including increased γH2AX foci, sister chromatid exchanges, micronuclei, and mitotic bridges that were not seen in Parp1 null cells (Figure 5D–F and Supplemental Figure 7C). These abnormalities were only modestly increased in *Parp1/2^-/-^* cells. Alkaline comet assays further demonstrated a significant increase in DNA strand breaks in *Parp1^EA/-^* cells (Figure 5G). Following methyl methanesulfonate (MMS) treatment, DNA damage accumulated rapidly in *Parp1^EA/-^* cells, reaching maximal levels within 10 minutes, whereas *Parp1/2^-/-^* cells exhibited a slower accumulation of damage that peaked approximately 40 minutes after treatment, consistent with delayed processing of base excision repair intermediates. Collectively, these findings demonstrate that catalytically inactive PARP1, unlike complete loss of PARP1 and PARP2, profoundly disrupts DNA replication and the repair of base damage both during and outside of replication. The resulting accumulation of unresolved DNA repair intermediates arising from sister chromatid exchange and homologous recombination-dependent repair, replication associated DNA gaps, and genome instability provides a mechanistic basis for the severe hematopoietic toxicity induced by clinical PARP inhibitors.

### Inactive PARP1 also compromises the recruitment of PARP2

An apparent paradox raised by our findings is why niraparib-induced anemia is dependent on PARP1 but not PARP2, even though niraparib directly traps PARP2 (24) and mice expressing catalytically inactive PARP2 (*Parp2^E534A/E534A^*) develop lethal fetal anemia (40). In this context, we previously reported that PARP1 and its activity promote the initial/early recruitment of PARP2 to DNA damage sites via XRCC1 (41). Specifically, the BRCT1 domain of XRCC1 binds to PAR generated by PARP1, while the BRCT2 domain of XRCC1 interacts with PARP2 (41). Consistent with this model, loss of PARP1 catalytic activity or XRCC1 reduced early PARP2 recruitment by approximately 50% (Figure 5H). Consequently, pharmacological inhibition of PARP1 suppresses PAR synthesis and simultaneously limits the amount of PARP2 that can be recruited and subsequently trapped at sites of DNA damage. We propose that this reduction in PARP2 recruitment by 50% keeps PARP2 trapping below the threshold required to disrupt Okazaki fragment maturation and cause lethal anemia (Figure 5I). Consistent with this interpretation, mice expressing inactive Parp2 from one allele (*Parp2^E534A/-^*) survive to adulthood with only moderate anemia in contrast to lethal anemia of *Parp2^E534A/E534A^* mice (40, 42). This model also explains the delayed accumulation of PARP2 at DNA damage sites following PARP inhibition (40, 41, 43) and why genetic loss of PARP1, but not PARP2, profoundly reduces the sensitivity of human cancer cells to dual PARP inhibitors (21). Together, these findings indicate that in addition to physically blocking repair factors’ access to DNA lesions, inactive PARP1 also attenuates the recruitment of PARP2 to suppress hematopoiesis.

### Discussions

Our results establish that the presence of catalytically inactive PARP1, not PARP2 inhibition or reduced PARylation, is the primary driver of PARPi-associated hematopoietic toxicity *in vivo* (Figure 5I). The hypersensitivity of cloning expanding lymphocytes and erythropoiesis to PARPi, at least in part, reflects proliferation-associated toxicity and the unique differentiation-coupled rapid DNA replication in erythroblasts characterized by fast fork progression (44) that renders erythroblasts especially vulnerable to replication stress. In this context, loss of LIG1 implicated in Okazaki fragment maturation is well tolerated in other tissues/cells, but not erythroblasts (40, 42), presumably because the Lig3-dependent Okazaki fragment joining is unable to compensate for the loss of the major Lig1-dependent pathway in erythroblasts (40, 42). Because the same replication-associated toxicity underlies the antitumor efficacy of PARPi, PARP1-selective inhibition is unlikely to eliminate hematologic toxicity. Indeed, the PARP1-selective inhibitor saruparib produced a similar spectrum of PARP1-dependent hematologic toxicity as niraparib, which was fully rescued by PARP1 deletion (Figure 3). Moreover, feedback from anemia (EPO) and thrombocytopenia (TPO) to HSPCs may explain how impaired erythropoiesis and megakaryopoiesis promote PARPi-associated clonal hematopoiesis(45). Together, these findings support inactive PARP1, generated genetically or pharmacologically, as a major driver of replication-associated PARPi hematologic toxicity.

Mechanistically, we showed that inactive PARP1, either via pharmacological inactivation or genetic mutation, physically blocks the repair and processing of diverse DNA lesions, including but not limited to ligase-mediated nick ligation, polymerase-mediated 3′-OH extension, as well as TDP1-mediated processing of Topo1-cc. Moreover, we demonstrated that the severe genomic instability (e.g., mitotic bridges, SCE) associated with PARP inhibition in HR proficient cells also arises from the presence of inactive PARP1 rather than loss of PARP catalytic activity. Cells expressing inactive PARP1, but not PARP1&2 double-deficient cells, displayed impaired replication fork progression and single-strand break repair, leading to increased mitotic breaks, elevated sister chromatid exchanges, and accumulation of S1 nuclease-sensitive gaps. These findings provide a coherent link between the molecular mechanism, cellular phenotype, and physiological consequence of PARP1 trapping.

So, if PARP1 selective inhibitors are not able to fully mitigate the hematological toxicity associated with PARP inhibition, what would be the solution? Beyond careful patient monitoring and intermittent dosing to allow BM recovery, we showed that CDK4/6 inhibition reduced *Parp1^Δ/E988A^*cell death (Figure 5B). CDK4/6i have recently been reported to mitigated clonal hematopoiesis in patients exposed to genotoxic chemotherapy (45), representing a promising strategy to limit PARPi hematological toxicity. Alternatively, non-trapping PARP inhibitors might be another option if therapeutic effects can be preserved, particularly when used in combination with other DNA damage agents.

## Materials and Methods

### Mice and *in vivo* treatment

#### Sex as a biological variable

Our study examined male and female animals, and similar findings are reported for both sexes.

#### Generation of the Parp1 conditional knockout mouse model

To generate the Parp1 conditional allele, a targeting vector was constructed to flank the exon 4 of murine PARP1 with loxP sites using homology targeting. Briefly, a neomycin selection cassette flanked by FRT and a single loxP site was inserted at intron 3, and another loxP site was inserted in intron 4 (Figure S2A). The targeting construct was linearized with SmaI and electroporated into the mouse embryonic stem (ES) cells (129/Sv background) and Neo-resistant clones were selected with G418. Correctly targeted clones were identified by Southern blot analysis following ScaI digestion and hybridization with a 3′ probe located outside the homology arms (Figure S2B). The probe was amplified by PCR using the following primers: 5′-GAACCCGGTGAAGTACCAAG-3′ & 5′-CAGTACTGATATGAGAAAGGGTGAGA-3′. The expected fragment sizes were 8.2 kb for the germline allele and 5.5 kb for the targeted allele. The successfully targeted clones were validated via southern blot analysis with a NeoR probe for single integration and sequenced to verify the loxP sites before being injected for germline transmission. The chimeric mice were subsequently crossed with constitutively FLPase-expressing *Rosa26a^Flp/Flp^* mice (46) (Jax stock number: 003946, also in 129/Sv background) to remove the NeoR cassette and create the conditional lines targeting Parp1.

#### Other mouse strains and genetic models

*Parp1^-/-^* mice were described previously and generously provided by Dr. Zhaoqi Wang (10). *Parp2^-/-^* and Parp1-E988A-Neo mice were generated and described previously (29, 30). Due to the germline transmission defects of *Parp1^⁺/E988A^* mice (30), somatic expression of the catalytically inactive Parp1-E988A allele was achieved by crossing *Rosa26a^Flp/Flp^;Parp1^flox/flox^* mice with *Rosa26a^Cre-ERT2/Cre-ERT2^;Parp1^E988A-neo/+^* mice (the E988A-neo allele contains a FRT-flanked Neo) to generate the *Rosa26a^Flp/Cre-ERT2^;Parp1^E988A-neo/flox^* mice and control *Rosa26a^Flp/Cre-ERT2^;Parp1^+/flox^* mice. Among them, the *Rosa26a^Flp^* allele expresses FLP constitutively, including in germline (46) and the *Rosa26a^Cre-ERT2^* contains an estrogen fused Cre-recombinase that requires tamoxifen/4OHT for nuclear translocation (33).

*Rosa26a^+/Cre-ERT2^;Parp1^flox/flox^;Parp2^flox/flox^* were generated to somatically delete both Parp1 and Parp2. The Parp2^flox^ allele was generously provided by Dr. Dantzer (34). Cre-recombination was induced by oral gavage of tamoxifen (100 mg/day in sunflower oil) for two consecutive days. Mice were analyzed 10 days after tamoxifen administration. Successful expression or deletion in bone marrow, spleen, and thymus was verified by PCR genotyping using genomic DNA extracted from these tissues. Genotyping for Parp1-E988A was performed as detailed before (30). Genotyping primers for Parp1-flox: 5′-CAGTCATGCGAATTGTACACC-3′, 5′-CAGCGGTCAATCATACCCAG-3′, and 5′-TGCTCCCCAGATCTCCACGA-3′ (WT allele, 280 bp; floxed allele, 370 bp; KO allele, 600 bp). Genotyping primers for Parp2-flox: 5′-CCCCAAACCAGAGTCCCATCC-3′ and 5′-CTCGAGTGTTTCACTGTGAGGGAG-3′ (WT allele, 494 bp; floxed allele, 657 bp; KO allele, 266 bp). Both male and female mice were used. Unless otherwise indicated, adult mice (6-12 weeks of age) were analyzed. At least three mice per genotype (typically >6) from independent litters were used. Sample sizes were determined via power calculations. Experiments were not randomized, and investigators were not blinded.

### In vivo PARP inhibitor treatment

To induce acute hematopoietic toxicity, young adult mice with indicated genotypes were orally gavaged daily with indicated PARP inhibitors for 5 consecutive days (niraparib, 60 mg/kg; saruparib, 12 mg/kg in 0.5% methylcellulose solution with 20% of DMSO as a control). Complete blood counts were performed by the Institute of Comparative Medicine at Columbia University using a Genesis analyzer (Oxford Science Inc.).

### Primary cell line derivation and culture

The MEFs were derived from E13.5–E14.5 embryos using standard procedures (47, 48), and immortalized via infection with retrovirus carrying SV40 large and small antigens (35, 49). Due to germline transmission defects, the *Rosa26a^Flp/Cre-ERT2^;Parp1^+/f^*and *Rosa26a^Flp/Cre-ERT2^;Parp1^EA/f^* MEFs were derived from E13.5-E14.5 embryos obtained from the cross between *Rosa26a^Cre-ERT2/Cre-ERT2^;Parp1^+/E988A-neo^*and *Rosa26a^Flp/Flp^;Parp1^flox/flox^* parents (30). The v-Abl kinase-transformed B cell lines (v-Abl B) were derived as previously described (47, 50–53). Briefly, single-cell bone marrow suspensions from <4 week-old mice of the indicated genotypes were infected with retrovirus encoding p120 minimal v-Abl kinase (54) and expanded for 6–8 weeks until stable clonal outgrowth was established. MEFs and v-Abl pre-B cells were cultured in DMEM (Gibco, 12430-062) supplemented with 15% FBS (Hyclone, SH30071.03), 1× MEM non-essential amino acids (Gibco, 11140-050), 1 mM sodium pyruvate (Gibco, 11360-070), 2 mM L-glutamine (Gibco, 25030-081), 120 μM β-mercaptoethanol (Fisher, 03446I-100), and 50 U/mL penicillin/streptomycin (Gibco, 15140122).

The *Rosa26a^Flp/Cre-ERT2^;Parp1^+/f^* and *Rosa26a^Flp/Cre-ERT2^;Parp1^EA/f^* ES cells were derived at the HICCC Transgenic Mouse Core (Columbia University) by superovulating ∼3-week-old *Rosa26a^Flp/Flp^;Parp1^flox/flox^* female mice mated with adult *Rosa26a^Cre-ERT2/Cre-ERT2^;Parp1^+/E988A-neo^*males. E3.5 blastocysts were cultured on irradiated (30 Gy) fibroblast feeders in DMEM (Gibco, 12430-062) supplemented with 20% FBS (Hyclone, SH30396.03), 1× MEM non-essential amino acids (Gibco, 11140-050), 1 mM sodium pyruvate (Gibco, 11360-070), 2 mM L-glutamine (Gibco, 25030-081), 120 μM β-mercaptoethanol (Fisher, 03446I-100), 50 U/mL penicillin/streptomycin (Gibco, 15070-63), and LIF (provided by the core facility) until inner cell masses attached and expanded.

### Flow cytometry analyses of hematopoietic cells

Flow cytometry analyses were performed as previously described with minor modifications (29, 40, 55). Single-cell suspensions were prepared from bone marrow, thymus, and spleen. Splenocytes were incubated in ACK lysis buffer (Lonza, BP10-548E) for 5 min at room temperature prior to processing. For B lymphocyte analysis, BM and spleen cells were stained with a cocktail of directly conjugated antibodies including FITC anti-mouse CD43 (BD Pharmingen, 553270), PE anti-mouse IgM (SouthernBiotech, 1020-09), PE/Cyanines5 anti-Hu/Mo CD45R (B220) (eBioScience; 15-0452-83), and APC anti-mouse TER119 (BioLegend, 116211). For T lymphocyte analysis, BM and spleen cells were stained with PE anti-mouse CD4 (BioLegend, 557308), FITC anti-mouse CD8a (BioLegend, 100706), PE/Cyanines5 anti-mouse CD3ε (eBioscience, 15-0031-83), and APC anti-mouse TCRβ (BD Pharmingen, 553174). For myeloid and erythroid cell analysis, BM cells were stained with FITC anti-mouse/human CD44 (BioLegend, 103006), PE anti-mouse CD71 (BioLegend, 113807), PerCP/Cyanine5.5 anti-mouse CD117 (c-Kit) (BioLegend, 105823), PE/Cyanine7 anti-mouse/human CD45R/B220 (BioLegend, 103222), APC anti-mouse Ly6G/Ly6C (Thermo Fisher Scientific, 17-5931-81), APC-eFluor™ 780 anti-mouse TER119 (Thermo Fisher Scientific, 47-5921-82), Alexa Fluor 700 anti-mouse CD11b (Thermo Fisher Scientific, 56-0112-82), Pacific Blue™ anti-mouse CD90.2 (Thy1.2) (BioLegend, 140306) and Brilliant Violet 510™ anti-mouse CD41 (BioLegend, 133923) for 10 min at room temperature (RT). For HSPC analysis, BM cells were first labeled with a cocktail of biotin-conjugated lineage-specific antibodies including anti-mouse TER119 (BioLegend, 116204), anti-mouse CD4 (BioLegend, 100404), anti-mouse CD8a (BioLegend, 100704), anti-mouse CD3ε (BioLegend, 100304), anti-mouse CD5 (BD Bioscience, 553018), anti-mouse/human CD45R/B220 (BioLegend, 103204) and anti-mouse Ly-6G/Ly-6C (Gr-1) (BioLegend, 108404) at 4 °C for 1 h. Then cells were washed and incubated with a cocktail of fluorophore-conjugated antibodies including FITC anti-mouse CD34 (Thermo Fisher Scientific, 11-0341-82), PE anti-mouse Ly-6A/E (Sca-1) (BioLegend, 108108), PE/Cyanine7 anti-mouse CD16/32 (BioLegend, 101318), APC anti-mouse CD117 (c-kit) (BioLegend, 135108), APC/Cyanine7 Streptavidin (BioLegend, 405208) and Brilliant Violet 510 anti-mouse CD41 (BioLegend, 133923) at RT for 10 min. The flow cytometry data were collected on either LSR II (BD) with BD FACSDiva software or Attune NxT (Thermo Fisher Scientific) with software v4.2. All flow cytometry data were analyzed using FlowJo V10.

### Western blotting

Western blotting was performed as detailed before (21, 30). Briefly, whole-cell lysates were prepared in modified RIPA buffer (50 mM Tris-HCl pH 7.4, 150 mM NaCl, 1% Triton X-100, 0.5% sodium deoxycholate, 0.1% SDS, 1 mM EDTA, 10 mM NaF) supplemented with 2 mM PMSF, 1× protease inhibitor cocktail (Roche, 11697498001), and 0.5 μL/mL Benzonase Nuclease HC (EMD Millipore, 71205-25KUN). Proteins were resolved by SDS-PAGE, transferred to PVDF membranes, and immunoblotted with antibodies against α-tubulin (Calbiochem, CP06), PAR (R&D, 4335-MC-100), and PARP1 (Cell Signaling Technology, 9542S).

### Sister chromatid exchange

Sister chromatid exchange assays were carried out with sequential labeling using classical protocols (56, 57). Briefly, *Rosa26a^Flp/Cre-ERT2^;Parp1^+/f^* and *Rosa26a^Flp/Cre-ERT2^;Parp1^EA/f^*ES cells at ∼passage 9 were seeded on feeder-free 60 mm dishes and treated with 200 nM 4-hydroxytamoxifen (Sigma, H7904) for 24 h, then 10 μM BrdU (Sigma, B9285) for another 24 h. Colcemid (0.3 μg/mL KaryoMax; Gibco, 15212-012) was added for 40 min, after which cells were trypsinized, swelled in 0.075 M KCl for 20 min, and fixed in methanol:acetic acid (3:1) for 10 min; centrifugation (700 rpm, 4 °C, 10 min) and fixation were repeated twice. Cells were resuspended in 500 μL fixative and dropped onto slides, which were steamed over boiling water, air-dried, and inspected by microscopy to confirm adequate metaphase spreads. After aging ≥24 h, slides were stained in 0.5× SSC containing 2 μg/mL Hoechst 33258 (Sigma, 14530) for 45 min and exposed to UV (Stratagene Stratalinker 2400, ∼40 kJ) for 40 min, washed in 0.5× SSC at 60 °C, air-dried, and mounted in DAPI-containing antifade medium (VECTASHIELD, H-1200). Metaphase images were acquired on a Zeiss Imager Z2 with Metafer 4, and 50–100 metaphases per genotype were scored.

### Alkaline comet assay

The v-Abl transformed *Rosa26a^Flp/Cre-ERT2^;Parp1^EA/f^* and *Rosa26^+/Cre-ERT2^;Parp1^flox/flox^;Parp2^flox/flox^*pre-B cells were derived from mice (<4 weeks) of the corresponding genotypes, cultured >2 months, and genotype-validated prior to experiments (35, 53, 58). Conditional alleles were deleted/recombined by treatment with 200 nM 4-hydroxytamoxifen for 48 h; WT pre-B cells were processed in parallel. Cells were then exposed to 0.1 mg/mL MMS for 10, 20, or 40 min, harvested simultaneously, and analyzed by alkaline comet assay per the manufacturer’s instructions (Trevigen). Images were acquired on a Nikon 80i fluorescence microscope (Plan Apo 20×/0.75 objective) and tail moments quantified with CometScore (v1.5).

### DNA fiber and S1 nuclease sensitivity assay

The DNA fiber assay and S1 nuclease sensitivity were measured as detailed before (59) with minor modifications. Briefly, *Rosa26a^Flp/Cre-ERT2^;Parp1^EA/f^* and *Rosa26^+/Cre-ERT2^;Parp1^flox/flox^;Parp2^flox/flox^*cells were treated with 200 nM 4-hydroxytamoxifen for 48 h, then pulse-labeled sequentially with 50 μM CldU (Sigma, C6891) for 20 min followed by 250 μM IdU (Sigma) for 60 min. Cells were harvested, washed in ice-cold PBS, and resuspended in lysis buffer (0.5% SDS, 200 mM Tris-HCl pH 7.4, 50 mM EDTA). Cell suspensions were spread by gravity onto Superfrost slides, air-dried, fixed in methanol:acetic acid (3:1) for 5 min, rehydrated in PBS for 10 min, and denatured in 2.5 M HCl for 30 min at room temperature. Slides were blocked in PBS containing 0.1% Triton X-100 and 5% BSA for 1 h, incubated with rat anti-BrdU (1:100, Abcam ab6326, recognizing CldU) and mouse anti-BrdU (1:100, BD Biosciences 347580, recognizing IdU) for 1.5 h, then with goat anti-rat Alexa Fluor 594 and goat anti-mouse Alexa Fluor 488 secondary antibodies (1:300; Thermo Fisher Scientific) for 45 min. Slides were mounted in antifade medium (Vectashield, H-1000-10) and dried overnight. Fibers were imaged on a Zeiss AxioImager Z2 (40×/0.75 objective) with the ISIS fluorescence imaging system and measured using ImageJ.

### Lig1-mediated nick ligation assay

Lig1 is purified as detailed before (40). LIG1-mediated nick ligation was assayed using a duplex containing a single ligatable nick assembled by a 5′ p^32^-labeled oligo (5′-CGC CAG GGT TCT GAG CAC AGT CAC GAC-3′) and an unlabeled oligo (5′-PO₄-AGT AAC ACG ACG GCC AGT GCT G-3′) with their complementary template (5′-CAG CAC TGG CCG TCG TGT TAC TGT CGT GAC TGT GCT CAG AAC CCT GGC G-3′). Purified PARP1 was preincubated with 0.5 pmol substrate in 20 mM HEPES-NaOH (pH 7.5), 100 mM NaCl, 10 mM MgCl₂, 2 mM ATP, and 0.1 mg/mL BSA, with or without 0.2 mM NAD⁺, at 37 °C for 15 min, followed by the addition of LIG1 (0.25 pmol) and further incubation at 37 °C for 5 min. Reactions were stopped with 2% SDS, 1× formamide loading buffer, and heat denaturation at 100 °C for 5 min. Products were resolved on 15% polyacrylamide/8 M urea gels (National Diagnostics; pre-run 2 h at 1400 V, run 1 h at 1200 V) and detected by phosphorimager analysis on a Typhoon FLA 7000 (GE Healthcare, 650 nm excitation) and quantified with ImageJ (v1.52a).

### Recombinant protein expression and purification of TDP1 and PARP1

An open reading frame containing human TDP1 or PARP1 was cloned in frame with an N-terminal TwinStrep-ZB-tag, followed by a TEV-protease cleavage site in a pNIC plasmid. PARP1 E988A was generated by introducing point mutation using the Q5 site-directed mutagenesis kit (New England BioLabs), following manufacturer’s instructions. Mutations were confirmed by Sanger sequencing. BL21 (DE3) Rosetta Escherichia coli cells were transformed with the corresponding plasmids for protein expression. Cells were grown in Terrific broth (TB) medium at 37°C to OD_600_ 0.7. Protein expression was induced by addition of 0.5 mM isopropyl-β-D-thiogalactoside (IPTG) for 16 h at 18°C. Cells were harvested, snap-frozen in liquid nitrogen and pellets were stored at - 80°C.

For TDP1 protein purification, cells were resuspended in lysis buffer (50 mM Tris/HCl pH 8, 500 mM KCl, 10 mM MgCl2, 0.1% IGEPAL, 0.04 mg/mL Pefabloc SC, complete EDTA-free protease inhibitor cocktail tablets, 1 mM tris(2-carboxyethyl)phosphine hydrochloride (TCEP)) and lysed by sonication. All subsequent steps were carried out at 4°C. Cell lysates were incubated with Benzonase nuclease (45 U/mL lysate) for 30 min, before cell debris was removed by centrifugation at 18,000 g for 30 min. Cleared and filtered supernatants were applied to Strep-Tactin^®^XT Superflow^®^ high capacity cartridges, equilibrated in wash buffer (50 mM Tris/HCl pH 8, 500 mM KCl, 1 mM TCEP) and protein was eluted in wash buffer with 50 mM biotin. Pooled fractions were applied on HiTrap Heparin HP affinity column and eluted in elution buffer (50 mM Tris/HCl pH 8, 1 M KCl, 1 mM TCEP). Protein was desalted in wash buffer and affinity tags were cleaved off over night with the addition of TEV protease at 1:10 mass ratio. Next, protein was loaded on a HiLoad® 16/600 Superdex® 200 pg column equilibrated in storage buffer (50 mM Tris/HCl pH 8, 150 mM KCl, 10% glycerol, 1 mM TCEP) for size exclusion chromatography. Eluted protein fractions were collected and concentrated with a 10 kDa molecular weight cut-off Amicon Ultra centrifugal filter. Concentrated protein was aliquoted, snap-frozen, and stored at -80°C.

For PARP1 protein purification, a similar protocol was followed with minor changes. Wash buffer used for the purification procedure contained 50 mM Tris/HCl pH 8, 300 mM KCl, 1 mM TCEP; and storage buffer contained 50 mM Tris/HCl pH 8, 250 mM KCl, 10% glycerol, 1 mM TCEP.

### T4, TdT and TDP1 dependent DNA enzymatic assays

Nicked or gapped duplex DNA substrates were generated by annealing fluorescently labeled oligonucleotides as follows: for the ligation assay, a 5′ Cy5-labeled oligo was annealed with a 5′-phosphorylated, 3′ FAM-labeled oligo to generate a nicked duplex; for the TdT assay, a 5′ Cy5-labeled oligo carrying a free 3′-OH was annealed with a 3′ FAM-labeled oligo to generate a gapped duplex with an exposed 3′-OH terminus; and for the TDP1 assay, a dual-labeled oligonucleotide carrying a 5′ Cy5 label and a 3′ FAM modification was annealed with a complementary scaffold strand to generate a nicked duplex bearing a 3′-phosphodiester-linked FAM adduct at the nick terminus. Increasing concentrations of purified PARP1^WT^ or PARP1^E988A^ (10, 25, 50, 75, 100 nM) were preincubated with 10 nM annealed substrate DNA in the presence or absence of NAD+ (2 mM) and Olaparib (10 μM) in 17.5 mM Tris-KCl (pH 8), 60 mM KCl, 2 mM MgCl₂, 0.1 mg/mL BSA, 5 mM TCEP and 3.5% Glycerol; with addition of 2 mM ATP for the T4 ligation reactions, or 0.25 mM CoCl_2_ and 0.2 mM dNTPs for the TdT extension reactions, at 30°C for 10 min. For the ligation reactions with T4 ligase, 5 μU T4 was then added, and the reactions were incubated further at 30°C for 30 min. For the 3′-OH extension reactions with TdT, 0.2 U TdT was then added, and reactions were incubated further at 37°C for 30 min. For the TDP1 3′-FAM cleavage reactions, 5 nM purified TDP1 was then added, and reactions were incubated further at 30°C for 30 min. All reactions were terminated by the addition of an equal volume of 2× denaturing stop solution (8M UREA, 15% Ficoll) followed by heating at 95°C for 10 min, and products were resolved by denaturing UREA–PAGE (12% acrylamide, 8 M urea, 1× TBE). Gels were imaged on a Bio-Rad ChemiDoc MP system using Cy5 and FAM fluorescence channels, and substrate consumption was quantified in ImageJ by measuring the fraction of unligated (ligation assay), unextended (TdT assay), or uncleaved FAM (TDP1 assay) signal relative to enzyme-free control reactions.

### Statistical analysis

Statistical analyses were performed using GraphPad Prism (v9). Data are presented as mean ± s.e.m. unless otherwise indicated. Statistical significance was determined using unpaired two-tailed Student’s t-tests unless otherwise specified. P < 0.05 was considered statistically significant. ns, p>0.05; * p<0.05; ** p<0.01; *** p<0.001.

### Study approval

All animal procedures were carried out in a specific pathogen-free facility at Columbia University Medical Center under protocols approved by the Institutional Animal Care and Use Committee (IACUC).

### Data availability

The data supporting the findings of this study are available within the paper and its Supplementary Information files. Further information and requests for resources and reagents should be directed to and will be fulfilled by the lead contact Dr. Shan Zha.

## AUTHOR CONTRIBUTION

X.L carried out all PARPi treatment experiments, generated and characterized the *Rosa26a^+/Cre-ERT2^;Parp1^flox/flox^;Parp2^flox/flox^* mice and the quantitative cell imaging analyses. Z.S. generated and characterized the *Rosa26a^Flp/Cre-ERT2^;Parp1^E988A-neo/flox^*mice and cells. W.J generated the mouse model carrying the Parp1 conditional allele with help from B.J.L. D.Y. in J.S. lab carried out the in vitro analyses of PARP1 and PARP1 E988A trapping with purified proteins with help from M.S.. D. M. performed the DNA fiber analyses. S.K.B in the A.E. T lab analyzed the impact of purified PARP1 on LIG1-mediated nick ligation. S.Z., X.L., and Z.S. planned the experiments and wrote the manuscripts. B.J.L. and F.Y helped with the breeding and colony management.

## ACKNOWLEDGMENT

We thank Dr. Alessandro Vindigni for advice on gap and DNA fiber analyses, Dr. Theresa Swayne at the HICCC Confocal Core for guidance on quantitative live-cell imaging experiments, and Dr. Victor Lin at the HICCC Transgenic Core for assistance with embryonic stem cell derivation. Due to space limits, we could not cite all original studies and apologize to colleagues whose important contributions were not referenced.

## Supplementary Figure Legends

**Supplemental Figure 1.**
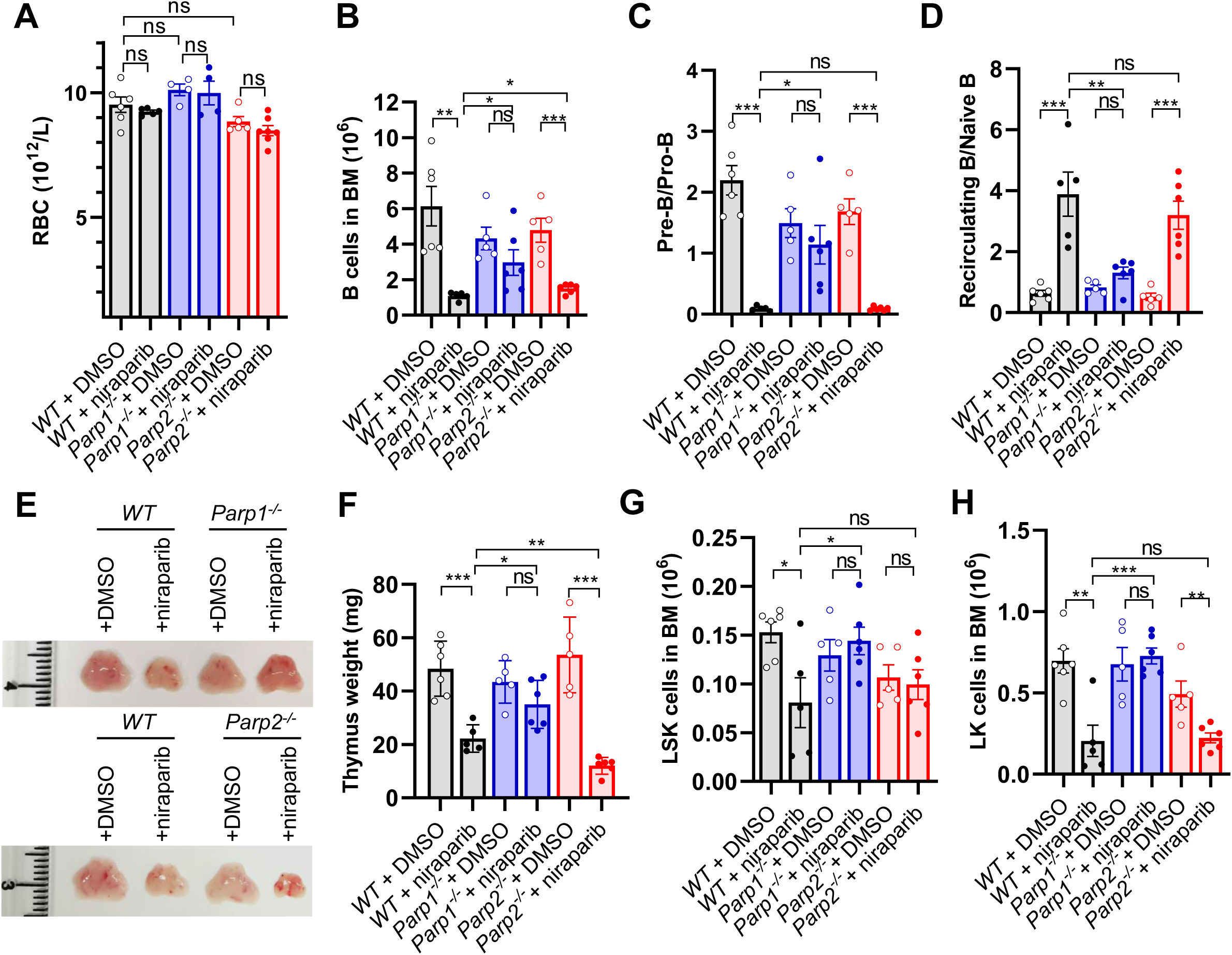
Extended analysis of *WT*, *Parp1^-/-^*, and *Parp2^-/-^* mice under niraparib treatment. (A) Red blood cells count in the peripheral blood from *WT*, *Parp1^-/-^* and *Parp2^-/-^* mice treated with niraparib. (B) Absolute number of BM cells from *WT*, *Parp1^-/-^* and *Parp2^-/-^*mice treated with niraparib. Ratios of (C)Pre-B/Pro-B cells and (D) Recirculating B/Naïve B cells in the BM from *WT*, *Parp1^-/-^* and *Parp2^-/-^* mice treated with niraparib. (E) Representative images of the thymus and (F) thymus weights from *WT*, *Parp1^-/-^* and *Parp2^-/-^* mice treated with niraparib. (G-H) Absolute numbers of (G) LSK and (H) LK cells in the BM from *WT*, *Parp1^-/-^* and *Parp2^-/-^* mice treated with niraparib.

**Supplemental Figure 2.**
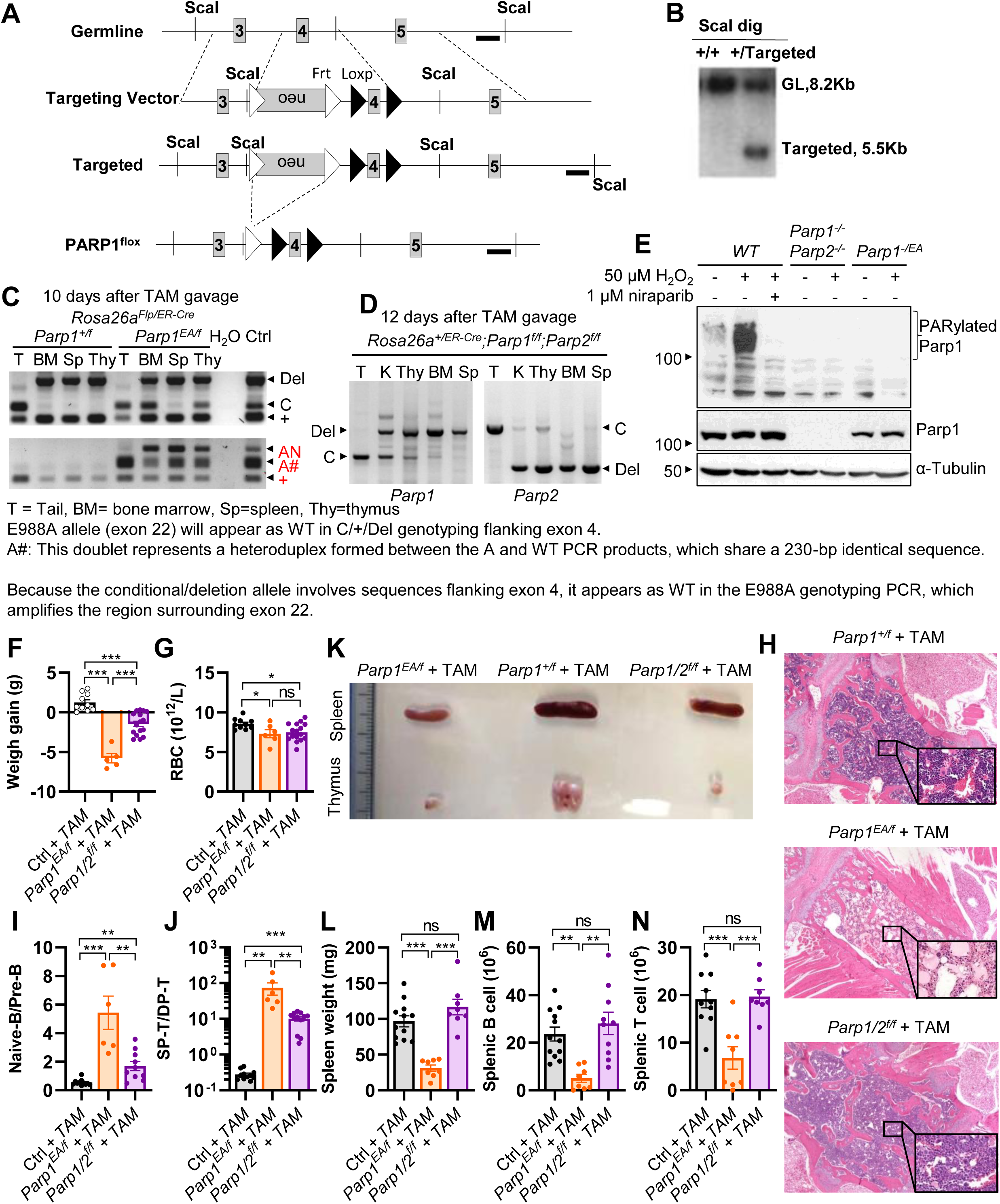
Generation of Parp1 floxed mice and extended characterization of somatic PARylation-deficient Parp1 and Parp1/2 DKO mice. (A) Targeting scheme of the *Parp1^flox^* allele. The exons are solid-filled grey boxes labeled with their corresponding numbers. FRT and loxP sites are indicated by open and black triangles, respectively. The probe used for Southern blotting is marked by a thick black line, and the positions of *SacI* restriction sites used for genomic DNA digestion are indicated. (B) Southern blot analyses of successfully targeted ES cells. Genomic DNA was digested with SacI restriction enzyme and detected by southern blot with the probe. (C and D) PCR results showing the recombination efficiency at FRT and loxP sites in the bone marrow, spleen, and thymus from *Rosa26a^Flp/Cre-ERT2^; Parp1^+/f^*, *Rosa26a^Flp/Cre-ERT2^;Parp1^EA/f^*and *Rosa26a^+/Cre-ERT2^;Parp1^f/f^;Parp2^f/f^*mice following tamoxifen gavage. Genomic DNA from tail tissue collected prior to gavage was used as a negative control for recombination. (E) Western blot showing the PARylation level in *WT*, *Parp1^-/-^Parp2^-/-^* and *Parp1^-/EA^* after following treatment with H₂O₂ and/or niraparib. (F) Body weight gain at day 10 of *Parp1^f/EA^*, *Parp1/2^f/f^* and control mice following tamoxifen gavage. (G) RBC counts in the peripheral blood from mice of the indicated genotypes. (H) Representative H&E stained BM sections from mice of the indicated genotypes. 40× images are shown, with black boxes indicating the regions displayed at higher magnification. Ratio of (I) naïve B/pre-B cells and (J) SP-T/DP-T cells from mice of the indicated genotypes. (K) Representative images of spleen and thymus from mice of the indicated genotypes. (L) Spleen weights, (M) splenic B cell numbers, and (N) splenic T cell numbers from mice of the indicated genotypes.

**Supplemental Figure 3.**
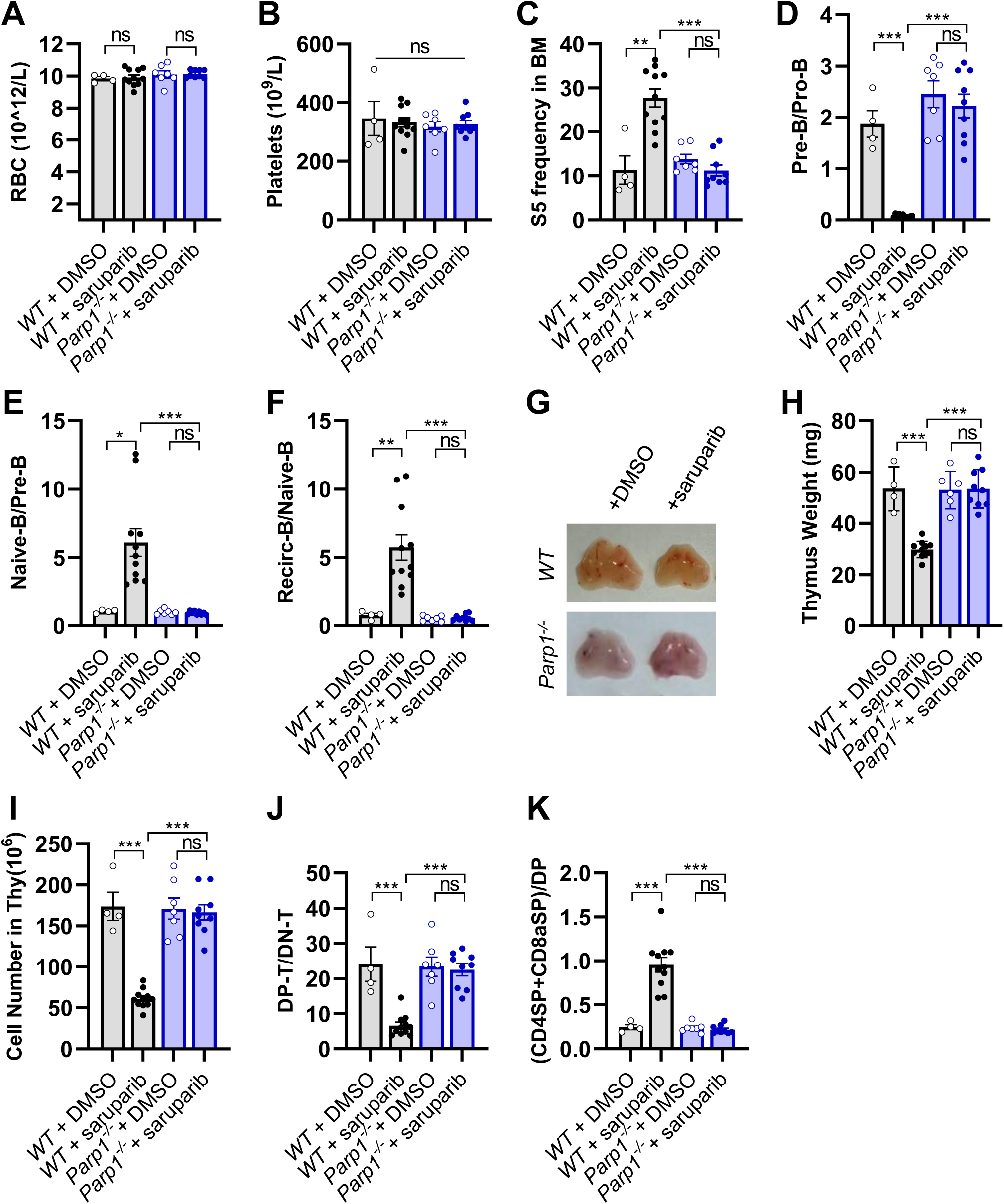
Extended analysis of *WT* and *Parp1^-/-^* mice under saruparib treatment. (A) RBC and (B) platelet counts in peripheral blood from *WT* and *Parp1^-/-^* mice treated with saruparib. (C) Frequency of S5 erythroid cells in the BM *WT* and *Parp1^-/-^* mice treated with saruparib. Ratios of (D) pre-B/pro-B, (E) naïve B/pre-B, and (F) recirculating B/naïve B cells in the BM from *WT* and *Parp1^-/-^* mice treated with saruparib. (G) Representative images, (H) weights, and (I) absolute cell number of the thymus from *WT* and *Parp1^-/-^* mice treated with saruparib. Ratios of (J) DP-T/DN-T and (K) SP-T/DP-T cells in the thymus from *WT* and *Parp1^-/-^* mice treated with saruparib.

**Supplemental Figure 4.**
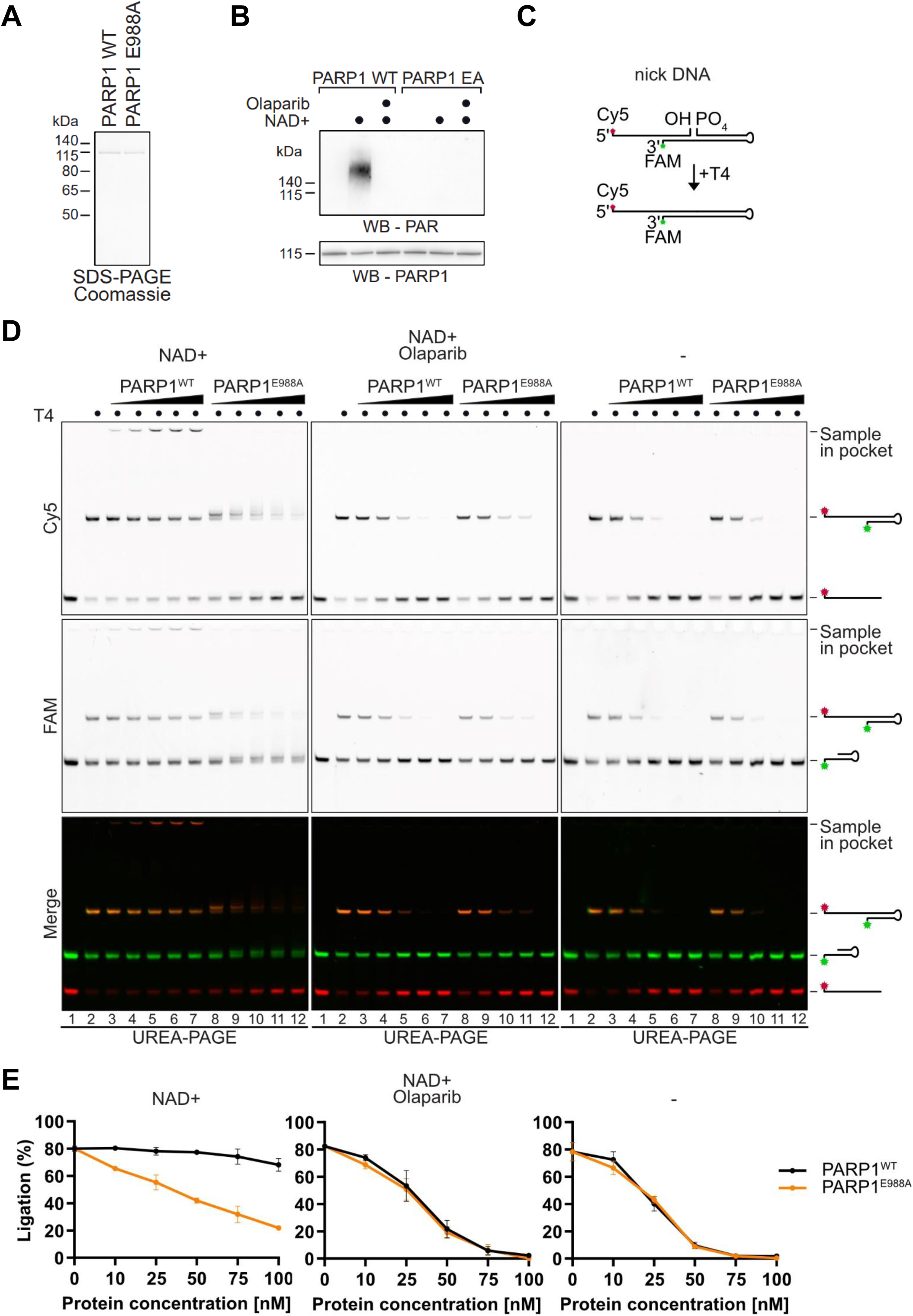
Inactive PARP1 blocks T4 DNA ligase-mediated ligation. (A) Coomassie-stained SDS-PAGE gel showing recombinant purified PARP1 WT and PARP1 E988A used for *in vitro* assays. (B) Western blot images showing the *in vitro* PARylation activity of PARP1 WT and PARP1 E988A in the presence of NAD+ and olaparib. (C) Schematic of the *in vitro* T4 DNA ligase-mediated ligation assay of 5′-phosphorylated nicked DNA. (D) Representative images of UREA-PAGE gels showing the results of the *in vitro* T4 DNA ligase-mediated ligation assay performed in the presence of increasing amounts of recombinant PARP1 (WT or catalytically inactive E988A), with or without NAD⁺ and olaparib, as indicated. (E) Quantification of ligation assay shown in (D). Values represent the means ± SD of three independent experiments.

**Supplemental Figure 5.**
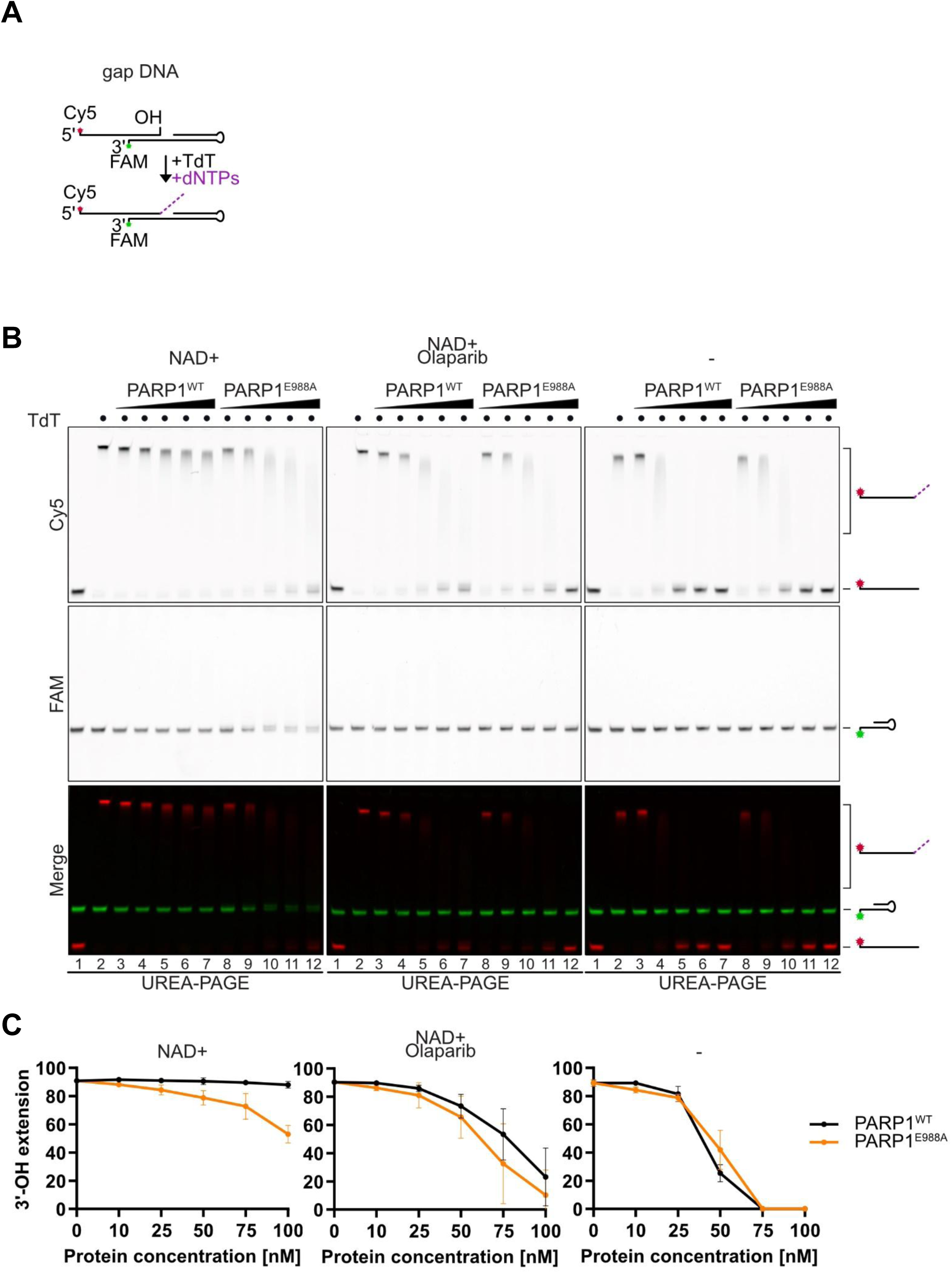
Inactive PARP1 blocks TdT-mediated DNA synthesis. (A) Schematic of the *in vitro* TdT-mediated DNA synthesis assay using 3′-OH gap DNA. (B) Representative images of UREA-PAGE gels showing the results of the *in vitro* TdT-mediated DNA synthesis assay performed in the presence of increasing amounts of recombinant PARP1 (WT or catalytically inactive E988A), with or without NAD⁺ and olaparib, as indicated. (C) Quantification of the DNA synthesis assay shown in (B). Values represent the means ± SD of three independent experiments.

**Supplemental Figure 6.**
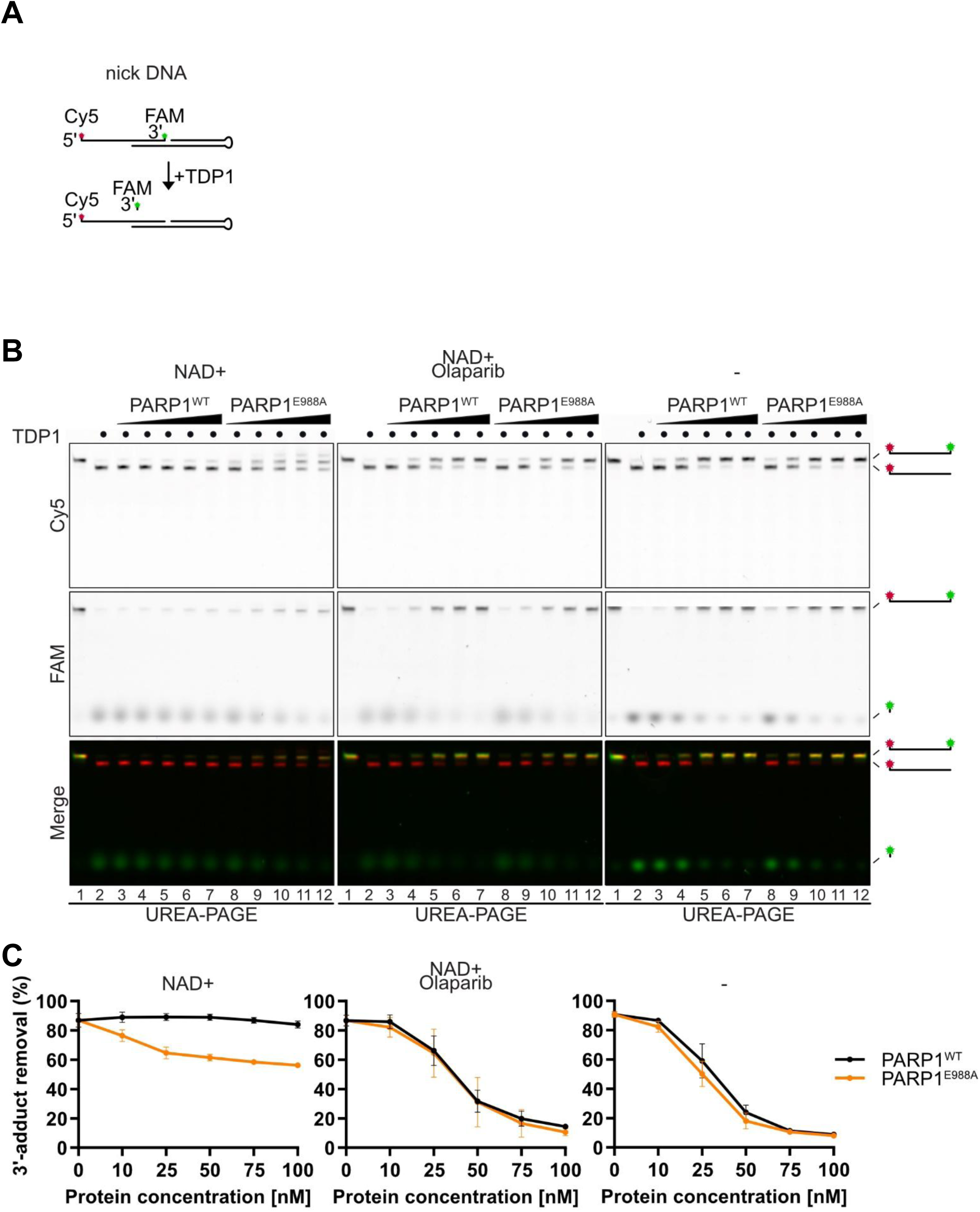
Inactive PARP1 blocks TDP1-mediated hydrolysis. (A) Schematic of the *in vitro* TDP1-mediated hydrolysis assay using 3′-phosphodiester-linked FAM-labeled nicked DNA. (B) Representative images of UREA-PAGE gels showing the results of the *in vitro* TDP1-mediated hydrolysis assay performed in the presence of increasing amounts of recombinant PARP1 (WT or catalytically inactive E988A), with or without NAD⁺ and olaparib, as indicated. (C) Quantification of hydrolysis assay shown in (B). Values represent the means ± SD of three independent experiments.

**Supplemental Figure 7.**
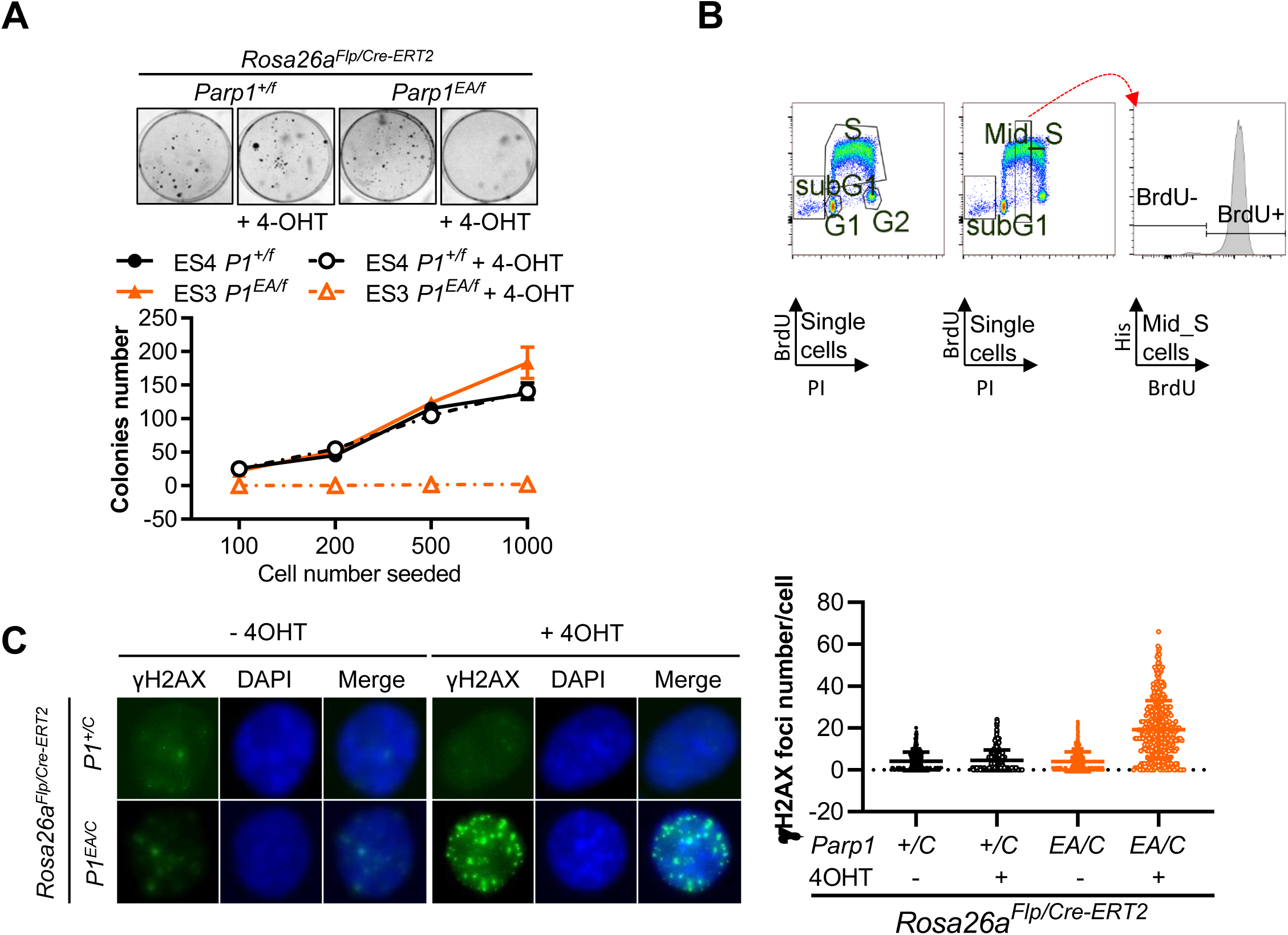
Inactive PARP1 impairs cell proliferation and induces genome instability. (A) Representative images and absolute numbers of colonies of ES cells derived from mice of the indicated genotypes. (B) Schematic of the gating strategy for cell cycle analysis in v-Abl B cells. (C) Representative immunofluorescence images and quantification of γH2AX foci v-Abl B cells of the indicated genotypes.

